# Interplay between human ribosomal proteins, PARP1, PARP2, HPF1 and histones

**DOI:** 10.1101/2025.09.15.676193

**Authors:** Andrey S. Krasnikov, Konstantin N. Naumenko, Mikhail M. Kutuzov, Yulia B. Zhakupova, Maxim O. Pavlov, Alexey A. Malygin, David Pastre, Dmitri M. Graifer, Olga I. Lavrik

## Abstract

ADP-ribosyl-transferases (ADP-ribose polymerases) PARP1 and PARP2 are critical players in DNA damage response in the nucleus. Being activated by a genotoxic stress, these enzymes utilize NAD^+^ to attach ADP-ribose chains to wide variety of proteins; ribosomal proteins (RPs) have been identified among the major targets of the modification in different cell lines. However, little remained known concerning the peculiarities of the reaction of RPs ADP-ribosylation itself. Here, we study ADP-ribosylation of human RPs within the large (60S) and small (40S) ribosomal subunits and those isolated from the subunits, with PARP1 and PARP2 *in vitro* using radioactively labeled NAD^+^. We fail to detect the modification of ribosome-bound RPs but observed ADP-ribosylation of certain ribosome-free RPs when we use total protein isolated from the subunits. RPs from the 60S subunit were globally more modified than those from the 40S subunit, and ADP-ribosylation of several 60S RPs (but not 40S) was considerably enhanced in the presence of histone PARylation factor 1 (HPF1). With all kind RPs, HPF1 switches the modification preferentially to their serine/tyrosine residues. Major targets of the 60S RPs ADP-ribosylation were identified as RPL4 (uL4), RPL6 (eL6) and RPL13A/RPL15 (uL13/eL15). The modification levels of particular RPs differently depend on the concentration of total RP; the most selective HPF1-dependent ADP-ribosylation occurs in RPL6 (eL6). When present simultaneously with histones, RPs win linker histone H1 in the competition for both PARPs; in contrast, core histones strongly compete with RPs for ADP-ribosylation. Possible functional assignments of ADP-ribosylation of RPs are discussed.

**Bullet points:**

- Free human ribosomal proteins are PARylated by PARP1 and PARP2;
- PARylation of ribosomal 60S proteins but not 40S ones is mostly HPF1-dependent;
- RPL4, RPL6 and RPL13A/RPL15 are the major targets of PARylation among 60S RPs;
- Linker histone H1 is a poor competitor to ribosomal proteins for PARPs;
- Core histones strongly competes with ribosomal proteins for PARPs.

## Introduction

ADP-ribosylation is a reversible posttranslational modification that can occur with a variety of proteins. Originally, it was found in the systems associated with cellular response to the DNA damage caused by different agents (reactive oxygen species, alkylating agents, UV-irradiation, etc.), but recent data indicate also implication of protein ADP-ribosylation in the regulation of many processes beyond DNA damage response (e.g., see [1-3]).

The reaction is catalyzed by enzymes from the family of ADP-ribosyl-transferases (ARTDs) called also poly ADP-ribose polymerases (PARPs). These enzymes use NAD^+^ as donor of ADP-ribose, and their family contains 17 members [4-5]. The most abundant member of the family is PARP1, which operates in the nucleus and is generally regarded to mainly contribute to poly (ADP-ribosyl)ation (PARylation) of nuclear proteins (e.g., see [6-7]). Another nuclear enzyme of this family, PARP2, catalyzes the synthesis of ADP-ribose and PARylation less efficiently than PARP1 [8-10]. Along with modification of various protein substrates, both enzymes can undergo automodification, and their activity is modulated by different proteins, e.g., by p53 or YB1 [1, 11-12]. Recently, an important co-factor of both PARP1 and PARP2, HPF1 (Histones PARylation Factor 1), operating in the case of PARylation of histones and autoPARylation, was discovered [13-14]. This protein, when forms a joint active center with PARP1 or PARP2, switches the preferential modification targets to serine residues [15-17]. At that, HPF1 can promote the initiation of ADP-ribosylation of histones or PARPs themselves and prevents the formation of long ADP-ribosyl chains, thereby enhancing the level of modified proteins carrying the shorter chains [17-18]. Role of HPF1 in the ADP-ribosylation of other substrates beyond PARP1/2 and histones is not yet studied. The majority of the PARP family members other than PARP1 and PARP2 typically operate in the cytosol (except PARP3 and PARP5a/b (TNKS1/2)) and attach, as a rule, only one ADP-ribose residue to proteins (mono ADP-ribosylation, MARylation) [19].

The spectra of ADP-ribosylated proteins (ADP-ribosylomes) identified in the cells treated with hydrogen peroxide, as an inducer of DNA damage, considerably depend on the cell type [6-7, 20]. At that, they can share a portion of common members including autoPARylated PARPs. Notably, many ribosomal proteins (RPs) are often present among ADP-ribosylated proteins too, and repertoire and/or abundance of RPs bearing such kind modification in some cell lines are rather large. Specifically, in MDA-MB-468 breast cancer cells about 20% of all ADP-ribosylated proteins are RPs [6], and in HeLa cells, 55 RPs of total known 80 ones have been found among ADP-ribosylated under genotoxic stress [20].

The presence of RPs among major targets for ADP-ribosylation implies that this modification might have somewhat relationships with biogenesis and/or functions of ribosomes, and, in the case of genotoxic stress, also with extra-ribosomal functions of these proteins. Such kind assumptions have been made even before identification of ADP-ribosylated RPs. So, in a *Drosophila* model system, almost 50% of nuclear poly(ADP-ribosyl)ated proteins along with PARP1 were localized in the nucleolus, where ribosome biogenesis occurs [21]. The above-mentioned data on ADP-ribosylomes obtained in other systems provide more ground to the assumption of a functional role of RPs ADP-ribosylation in the biogenesis of ribosomes and/or translation regulation as well as other functions associated to RPs. Notably, it has been reported that some amino acid residues identified as ADP-ribosylation targets in RPs are important players in the formation of the subunits structure or intersubunit bridges in the mature 80S ribosome [6]. Finally, the functional effects of specific MARylation of definite RPs with cytosolic PARPs within the ribosome have been recently described. Specific MARylation of several RPs from both ribosomal subunits (including RPL24 and RPS6) by PARP16 results in a decrease of the translation level and polysomes assembly in ovarian cancer cells, and this effect is not observed with normal fallopian cells [2]. Besides, specific MARylation of the 40S ribosomal subunit at the RP RACK1 by PARP14 reduces translation of specific mRNAs encoding crucial regulators of carcinogenesis and promotes formation of stress granules containing 40S RPs including RACK1 [3].

In spite of the above-mentioned data pointing to RPs as targets of the ADP-ribosylation in nucleus, there are more questions than responses in this field. Identification of RPs by mass-spectrometry analysis of peptides resulting from protein proteolysis, is mostly qualitative and does not enable the quantitative measure of preferences of PARPs towards specific RPs. It remains also unknown whether the modification occurs in ribosome-free RPs or in those within assembled ribosome subunits. Interrelations between ADP-ribosylation of RPs, histones (major components of the nucleus and nucleolus, which can compete with ribosomal proteins for PARPs) and PARP1 or PARP2 themselves are also obscure. Finally, the role of HPF1 in the modification of RPs is unknown; the data in this respect are limited, to our knowledge, to a single study [7], in which the dependence of recombinant RPS3A PARylation with PARP1 *in vitro* on HPF1 has been shown.

Here, to address to above issues, we carry out ADP-ribosylation of RPs using ^32^P-labeled NAD^+^ with PARP1 or PARP2 in the isolated human 40S and 60S ribosomal subunits and in the total ribosomal protein mixtures from these subunits. The subsequent electrophoretic analysis of the ^32^P-labeled PARylated proteins allows the quantitative comparison of the modification levels of particular RPs and the dependence of PARylation on HPF1 as well on the presence of histones as likely competition targets for PARylation in the nucleolus. We identify a marked preference for the modification 60S subunit RPs and reveal the dependence of ADP-ribosylation of particular 60S RPs (but not 40S ones) with PARP1 and PARP2 on the presence of HPF1. We find out that the 40S and 60S subunit RPs strongly inhibit PARylation of linker histone H1 likely in a non-competitive way, but they are weak competitors for PARPs with core histones. As well, we determine the dependence of the PARPs automodification level on the presence of the above-mentioned proteins.

## Materials and Methods

### Materials

The following materials and reagents were used in the work: N,N′-methylenebisacrylamide, tris(hydroxymethyl)aminomethane (Tris) (Amresco, USA); acrylamide, bisacrylamide (AppliChem, Germany); ammonium persulfate (APS) (PanReac, Spain); sodium dodecyl sulfate (SDS) (Fluka, Switzerland); nicotinamide adenine dinucleotide (NAD^+^), LB Broth (Sigma-Aldrich, USA); cells of the E. coli strain Rosetta(DE3) pLysS (Merck, USA); [α-32P]ATP with specific activity of 1000 Ci/mmol (Laboratory of Biotechnology, Institute of Chemical Biology and Fundamental Medicine, Siberian Branch of the Russian Academy of Sciences, Russia). Olaparib (AZD2281) was purchased from Sigma-Aldrich (cat. SML3705). The recombinant bovine poly(ADP-ribose) glycohydrolase (PARG) was kindly provided by E. Ilina (ICBFM, Novosibirsk, Russia). Core histones (the mixture of H2A, H2B, H3 and H4) were isolated from Gallus gallus erythrocytes and purified by Bio-Gel HT Hydroxyapatite chromatography (Bio-Rad, cat. #1300150) as described [22]. To obtain the mixture of core histones, after the chromatography the solution was dialysed against buffer (25 mM KH_2_PO_4_, pH 7.0, 0.5 M NaCl and 40% glycerol) [22]. Other reagents and buffer components were of extra pure and analytically pure grades.

### Proteins Expression and Purification

Histone H1 was overexpressed in Escherichia coli BL21(DE3) transformed with plasmid pLate31-H1 containing a coding sequence for human histone H1 and purified by Ni-NTA affinity chromatography (GE Healthcare, cat. GE17-5255-01), Mono-S chromatography (GE Healthcare, cat. GE17-5168-01) and ssDNA-cellulose chromatography (Sigma-Aldrich, cat. D8273).

Recombinant PARP1 and PARP2 were overexpressed in Sf9 insect cells and purified by 3-aminobenzamide Sepharose and Heparine Sepharose (GE Healthcare, United States) chromatographic steps as described earlier [23].

Recombinant HPF1 overexpressed in E. coli Rosetta (DE3)pLysS (Novogen, USA) and purified by Ni-NTA agarose (GE Healthcare, United States) affinity chromatography, Q-sepharose (GE Healthcare, United States) and Superdex 16/600 (GE Healthcare, United States) as described previously [17].

### Synthesis of radioactive NAD+

[^32^P]-Labeled NAD^+^ was synthesized in a reaction mixture containing 25 mM Tris-HCl (pH 7.5), 20 mM MgCl_2_, 2 mM β-nicotinamide mononucleotide, 1 mM ATP, 0.5 mCi [α-^32^P]ATP and 5 mg/mL nicotinamide mononucleotide-adenylyltransferase (NMNAT), for 1 h at 37°C. NMNAT was inactivated for 10 min at 65°C; the denatured enzyme was pelleted by centrifugation (12,000 g, 10 min at 4°C).

### Ribosomes and ribosomal proteins

60S and 40S ribosomal subunits were isolated from unfrozen human placenta and characterized as described [24]. Total ribosomal protein from the subunits was isolated using a standard procedure based on their acetic acid extraction from the subunits solution at a high Mg^2+^ concentration [25] with some alterations. At the final step, the proteins were precipitated by addition of 6 volumes of cold acetone, the pellet dissolved in 2 M urea up to a concentration of 5-25 µM and stored at -20°C.

### ADP-ribosylation of RPs

Catalyzed by PARP1 and PARP2 autopoly(ADP-ribosyl)ation of total ribosomal protein isolated from 60S and 40S ribosomal subunits, and those within intact 60S (40S) ribosomal subunits was carried out in standard 10 μl reaction mixtures containing 20 mM Tris-HCl, pH 7.5, 120 mM KCl, 10 mM MgCl_2_, 1 µM [^32^P]NAD^+^, 0.5 OD_260_/ml DNA-activator (dsDNA after incubation with DNase I), 0.5 μM PARP1 (PARP2), and 0.4-2.0 μM HPF1 (where indicated); in some experiments, the reaction mixtures as well contained histone H1 or core histones. The reaction was initiated by adding [^32^P]NAD^+^ to mixture preassembled on ice. After incubating the mixtures at 25°C for 20 min for PARP1 (PARP2), the reactions were terminated by the addition of SDS-PAGE sample buffer and heating for 5 min at 95°C. Where indicated, reaction mixtures were treated with PARG and olaparib (1 µM and 10 µM, respectively) and/or with 1 M hydroxylamine (NH_2_OH, pH 7.5) for 45 min at 37°C before addition of loading buffer. The reaction products were separated by 15% SDS-PAGE (a ratio between acrylamide and bis-acrylamide of 37.5:1); bands of proteins labelled with [^32^P]ADP-ribose were analyzed by using the Typhoon imaging system (GE Healthcare Life Sciences) and Quantity One Basic software (Bio-Rad). The radiolabelled signals of modified proteins were quantified as follows: the total (raw) signal of the smeared band of modified protein (indicated for each protein in the autoradiograms) was quantified and the same-size background signal of gel in the respective lane was subtracted from the raw signal. The quantitative data presented in histograms were obtained in at least three independent experiments.

### Identification of ADP-ribosylated RPs by MALDI-TOF mass-spectrometry

ADP-ribosylated RPs were resolved by SDS PAGE as described above, radioactive bands corresponding to the modified RPs were excised from the gel, subjected to trypsin digestion, the resulting peptides after their purification were analyzed by MALDI-TOF mass spectrometry as described [26] with minor alterations, and RPs were identified based on the Mascot database with the use of mMass software (more details are given in the Supplement 1).

## Results

### Ribosomal proteins undergo PARylation outside of the assembled ribosomal sbunits

First, we attempted to carry out PARylation of RPs with PARP1 and PARP2 in the mature human ribosomal subunits. Since it is known that the PARP activity is considerably regulated by damaged DNA, we used DNA from calf thymus treated with DNase I (hereinafter referred as the activated DNA or dsDNA activator) in our experiments. HPF1 was used as a cofactor of PARP1 and PARP2. However, we failed to detect a substantial amount of PARylated RPs within the ribosomal subunits (Figure 1a and 1b, lanes 5,6 and 9,10). On the other hand, with total protein isolated from the subunits, several RPs of the 60S ribosomal subunit (lanes 7 and 8) and of the 40S subunit (lanes 3 and 4) are able to be ^32^P-ADP-ribosylated in a reaction catalyzed by either PARP1 or PARP2 in the presence of activated DNA. The presence of HPF1 drastically changes the modification patterns of 60S RPs with both PARPs; one can see a considerable enhancement of the intensity of particular RP bands, which is more pronounced with PARP2. Noteworthy, this enhancement strongly varies for different proteins. Of three distinct bands of PARylated with PARP1 60S RPs present on the electrophoregrams (marked here and after as bands I, II and III, see Figure 1), intensity of the band I barely depends on the HPF1 presence. The intensity of the band III is moderately enhanced by HPF1, whereas the intensity of the band II is strongly enhanced (Figure 1a, compare lanes 7 and 8). With PARP2, the intensities of all RPs bands almost completely depend on the HPF1 presence (Figure 1b, lanes 7 and 8).

**Figure 1.**
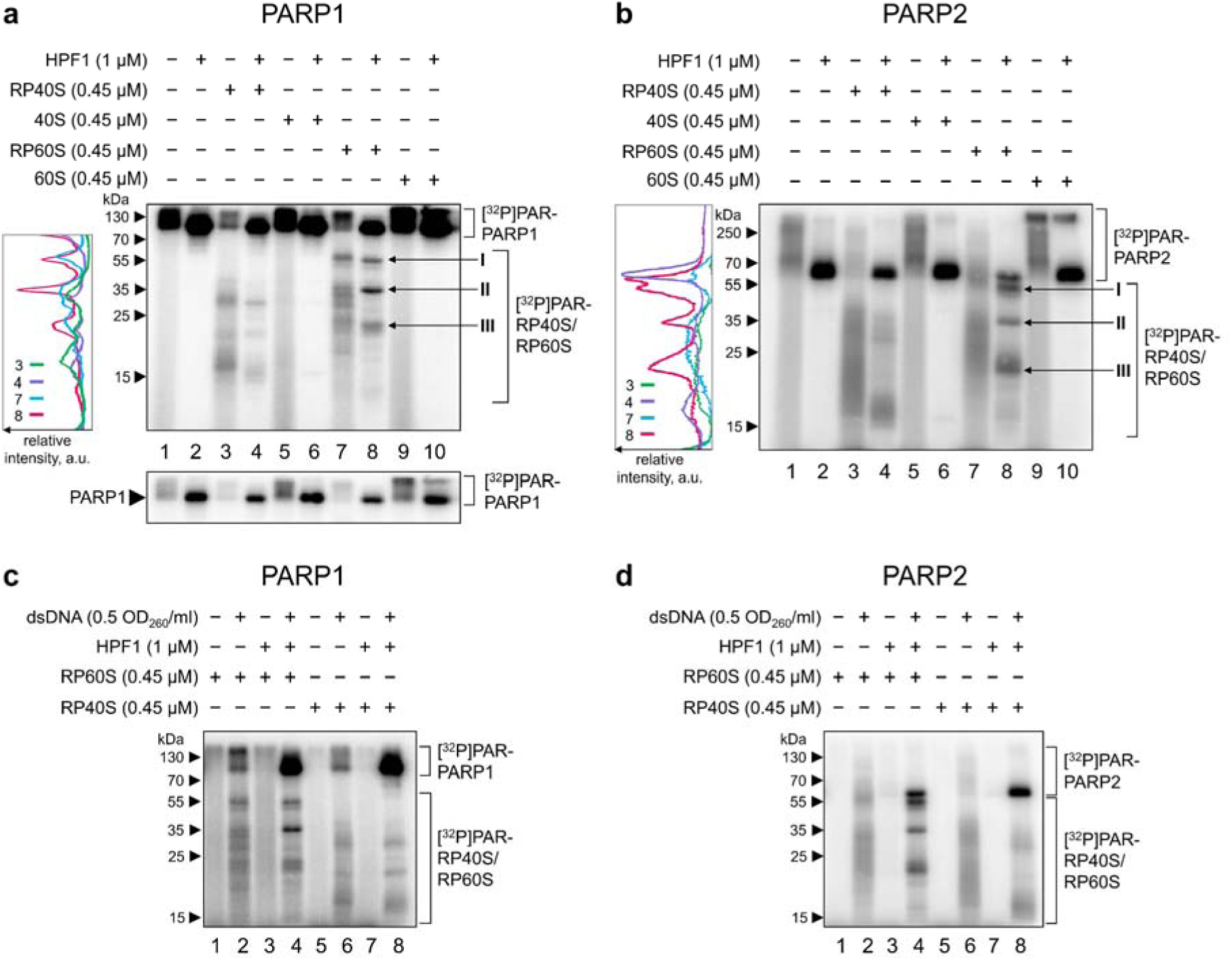
Dependence of PARylation of ribosomal proteins within and outside the ribosomal subunits with PARP1 and PARP2 on the presence of HPF1 and dsDNA-activator. Autoradiograms of the PARylation products separated by 15% SDS-PAGE. Covalent attachment of ^32^P-labelled ADP-ribose to 60S or 40S ribosomal proteins was performed by incubation of PARP1 (**a**) or PARP2 (**b**) with [^32^P]NAD^+^, dsDNA-activator, and with HPF1 (1□μM), where indicated. Lanes 3-4 and 7-8 correspond to the total protein isolated from the subunits, and lanes 5-6 and 9-10 to the proteins within the assembled subunits. Diagrams representing the distribution of the intensity among the bands are shown on the left of the autoradiograms (**a**) and (**b**). Positions of PARylated proteins and autoPARylated PARPs are indicated on the right (including three major bands I-III of the PARylated 60S subunit proteins); positions of the molecular weight markers are shown on the left. Top section of the autoradiograph of reduced exposure containing only the PARP1 signals is given below the main panel (**a**). Panels (**c**) and (**d**) show the dependence of PARylation of RPs with PARP1 and PARP2, respectively, on the presence or absence of the dsDNA-activator.

As one can see from the parallel experiments, the yield of the HPF1-dependent PARylation of 40S RPs is in general much less than that with 60S RPs (Figure 1a and b, compare lanes 4 and 7). Surprisingly, with the 40S subunit RPs, HPF1 does not enhance the intensities of the RP bands, and only makes them sharper (Figure 1a, b, lanes 3 and 4). As expected, PARylation of any kind RPs with both PARPs completely depends on the presence of DNA activator; without the latter, modification of RPs and the enzymes themselves is barely detectable both with and without HPF1 (Figure 1c, d).

Thus, several free ribosomal proteins are capable of ADP-ribosylation by PARP1 and PARP2 under certain conditions and the yield of PARylation of particular 60S ribosomal subunit RPs considerably depends on the presence of HPF1; the latter is not observed with the 40S subunit RPs. Ribosome-free RPs not only undergo PARylation, but also noticeably decrease the level of the PARPs autoPARylation both in the presence of HPF1 (Figure 1a, b, bottom panels, compare lanes 2 and 4 and lanes 6 and 8) and without the co-factor (compare lanes 1 and 3 and lanes 5 and 7). With PARP2 the effect is more explicit than with PARP1; intact ribosomal subunits do not substantially affect autoPARylation of both PARPs. For all that, as expected, in the presence of HPF1 the level of PARPs automodification is much stronger and the PAR chains are clearly shorter (the respective radioactive bands correspond to products with higher electrophoretic mobility). Similar observation as well could be done for automodification of PARP1.

### PARylation of certain RPs is HPF1-dependent

To explore the HPF1-dependency of the RPs modification more precisely, we examined the dependence of the ribosomal proteins PARylation with PARP1 and 2 on the concentrations of HPF1 and of the total RP. It turned out that the RPs modification patterns depend on the total RP concentration more significantly than on the HPF1 concentration (Figure 2). The dependence of the RPs’ PARylation pattern on the concentration of the total RP is especially pronounced with 60S RPs. So, of three major radioactive bands I-III of PARylated 60S RPs, only the middle one (II) appears as a major band already at the total RP concentration of 0.2 µM with both PARPs (Figure 2a and 2b). With PARP1, under these conditions, band III is also seen as a minor one; increase of the total RP concentration up to 0.45 µM leads to the appearance of the upper band, and further increase of the RP concentration up to 1.35 µM results in weakening of all three protein bands. With PARP2, both upper and lower bands appear in addition to the middle one at the 0.45 µM concentration, and at 1.35 µM RP the lower band significantly strengthens in contrast to the weakening other two ones. The effect of HPF1 concentration is not strong and may be the most reliably detected with PARP2 and smaller RPs corresponding to the lower band. In the latter case, at lower HPF1 concentration, one can see a trail above the major band, which disappears at higher concentrations of the co-factor. This is likely caused by the expected shortening of the PAR chains attached to the proteins upon the increase of HPF1 concentration [18]. The effect is better seen with smaller RPs (corresponding to the lower bands in each panel of the Figure 2) because the smaller protein, the larger contribution of the attached PAR chains to its electrophoretic mobility.

**Figure 2.**
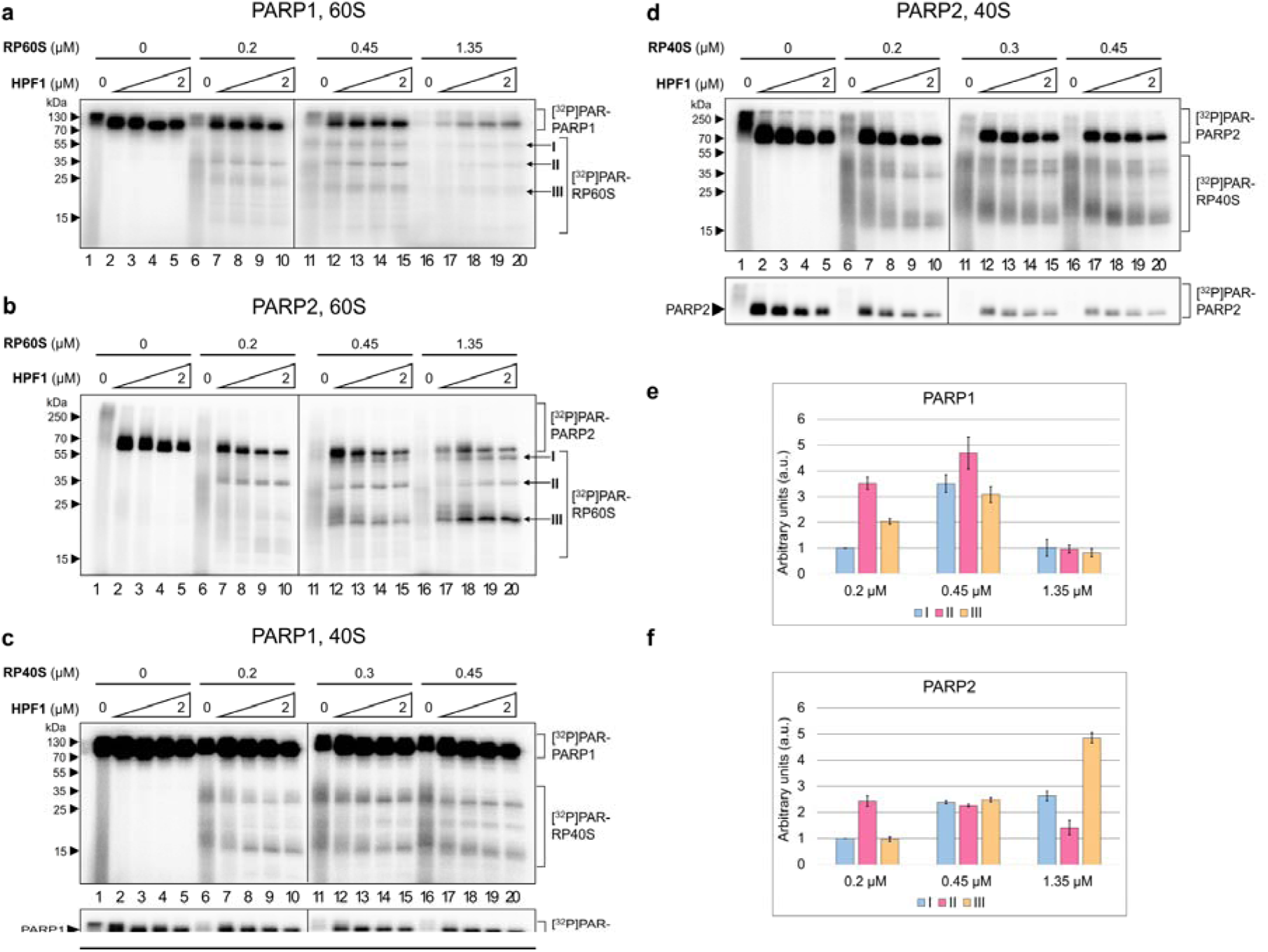
Dependence of the yield of PARylated RPs on the concentration of total RP and HPF1. Autoradiograms of the gels after separation of the products by 15% SDS-PAGE. Covalent attachement of ^32^P-labelled ADP-ribose to the total RP from the 60S (**a, b**) or 40S (**c, d**) subunits upon their incubation with PARP1 (**a, c**) or PARP2 (**b, d**) and [^32^P]-NAD^+^ in the presence of dsDNA-activator. Histograms (**e**) and (**f**) show relative intensities of the bands I, II and III assuming the intensity of the band I at 0.2 μM as 1 (the mean values calculated from three independent experiments). Increasing HPF1 concentrations used here were 0.4, 0.9, 1.5 and 2 µM. The positions of PARylated proteins and autoPARylated PARPs are indicated on the right sides of the autoradiograms; the positions of the molecular weight markers are on the left sides. Bands I-III are the same as in Figure 1. Parts of the autoradiographs with reduced exposure that contain only the PARP signals are given at the bottom.

Another feature of the HPF1-dependent RPs PARylation visible from the results presented in Figure 2 is that the sets of the modified bands with both PARPs are in general similar for the 60S RPs (bands I-III defined in Figure I a, b) but distinctly differ for the 40S ones. In the latter case, the modification with PARP2 seems to be less selective than with PARP1.

### HPF1 switches PARylation of both 60S and 40S RPs to serine residues

PARP1 and PARP2 have been described to mainly target ADP-ribosylation to Glu and Asp residues of substrate proteins [27]. HPF1 switches targets of ADP-ribosylation to Ser residues [14], and in some cases also to Tyr residues [7]. However, with the use of hydrolase ARH3 (ADP-ribosylhydrolase 3) that specifically cleaves the bond between ADP-ribose and Ser residues, it has been shown these residues are dominant targets for ADP-ribosylation by PARP1 and PARP2 in the presence of HPF1 [14]. Earlier studies demonstrated that hydroxylamine effectively removes ADP-ribose (ADPr) from Glu and Asp residues but does not remove ADPr from serines [14]. This confirms that ADPr is attached to Glu/Asp via ester bonds, which are labile to hydroxylamine, whereas Ser-ADPr (or Tyr-ADPr) likely involves a more stable ether linkage that resists cleavage under the same conditions [28]. To examine whether this is the case with the modification of RPs, we monitored here the stability of PARylated proteins to the treatment by hydroxylamine considering that the PAR modification of Ser/Tyr residues is stable under this treatment [29].

Indeed, the radioactive bands corresponding to ADP-ribosylated with PARP1 or PARP2 in the presence of HPF1 ribosome-free 60S subunit RPs do not considerably change their intensities after the hydroxylamine treatment (This is also true for the bands of the automodified PARPs, see Figure 3, compare lane 3 with lane 4 and lane 7 with lane 8). This implies that the ADP-ribosylated 60S RPs are mostly stable upon the hydroxylamine treatment, which is peculiar for proteins modified at Ser/Tyr residues. In parallel experiments without HPF1, the respective bands with hydroxylamine-treated samples practically disappear (lanes 2 and 6), as expected for proteins modified without HPF1, when targets of PARylation of proteins have to be Asp and Glu residues [14]. Similar regularities are observed with the 40S subunit RPs and both PARPs (see the respective lanes in Figure 3c, d). Thus, HPF1 switches mainly ADP-ribosylation of RPs to serine residues by a similar way as that of histones and PARPs.

**Figure 3.**
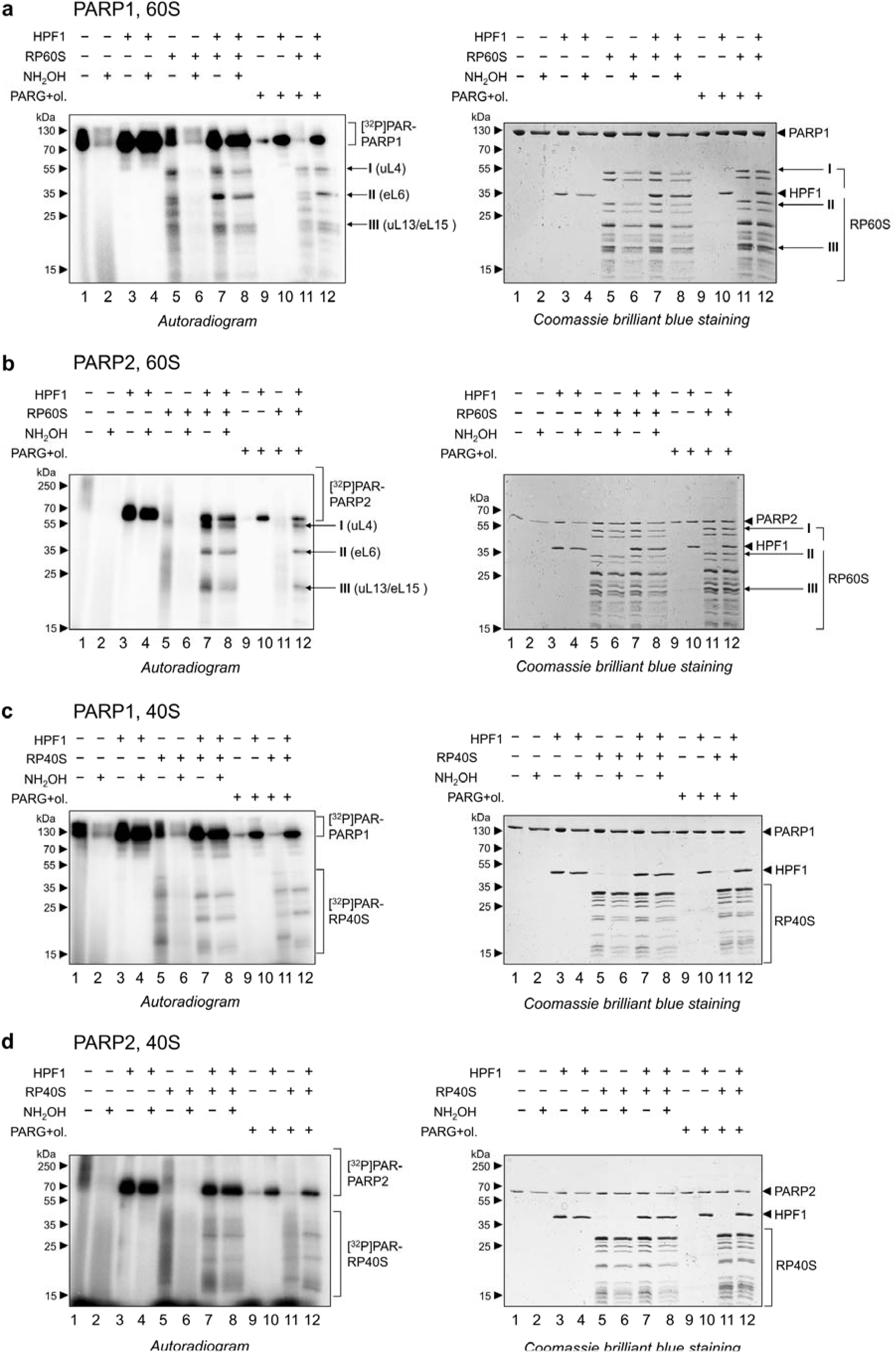
Examination of the HPF1-induced switch of the amino acid specificity of ADP-ribosylation of ribosomal proteins to Ser residues. Autoradiograms show covalent binding of ^32^P-labelled ADP-ribose to PARP1/PARP2 (0.5 μM) and 60S (**a, b**) or 40S (**c, d**) total ribosomal protein (0.45 μM) after incubation of the PARPs with [^32^P]NAD^+^ in the absence or presence of HPF1 (1 μM) with and without treatment with 1M hydroxylamine. The reaction mixtures also contained PARG (1 μM) and olaparib (10 μM), where indicated. Autoradiograms of the gels after separation of the products by 15% SDS-PAGE. On the right, Coomassie-stained gels corresponding to the autoradiograms are presented. It is seen that hydroxylamine treatment leads to some loss of the protein material in general. Therefore, we do not consider somewhat decrease of the intensity of the protein bands in lanes 8 as compared to lanes 7 as an indication for incomplete switch of the reaction. Bands I, II and III are the same as in Figures 1 and 2; RPs corresponding to these bands identified by mass-spectrometry (

Poly(ADP-ribosyl) chains of unknown length do not allow the identification of the modified peptides by mass-spectrometry. Therefore, to identify ADP-ribosylated RPs in the total mixture of 60S RPs, we treated proteins after the modification with poly ADP-ribose glycohydrolase (PARG) that hydrolyze PAR to a single ADP-ribose moiety attached to proteins [6-7, 20]. This treatment resulted in a substantial (approx. 2-fold; see Supplement 1, Figure S1) decrease in the intensities of radioactive bands corresponding to the modified 60S RPs with either PARP but did not cause a visible shift of the bands positions in the autoradiograms (Figure 3a and b, lanes 7 and 12). This implies that ADP-ribosyl chains attached to the 60S RPs are rather short. This suggestion is in agreement with a much more considerable decrease in the intensities of the bands corresponding to the PARP1/2 automodified in the absence of HPF1 (when the PAR chains are long [17]) after the treatment with PARG and olaparib, a PARP inhibitor (Figures 3a and b, lanes 1 and 9). Remarkably, with the 40S RPs ADP-ribosylated in the presence of HPF1, the PARG treatment leads only to a minor reduction of the intensities of the bands of the modified RPs (Figure 3,d, lanes 11 and 12) in contrast to those of the 60S RPs. This suggests that ADP-ribosyl chains attached to the 40S RPs are shorter than those attached to the 60S RPs and they most likely consist of 1-2 ADP-ribose residues.

60S RPs modified with PARP1 corresponding to three major radioactive bands I-III (Figure 1a, b) were identified by MALDI-TOF mass-spectrometry analysis. For this, the 60S RPs modified by PARP1 and treated with PARG were separated by SDS-PAGE and proteins corresponding to the above bands were digested with trypsin (see Supplement 1, Figure S2 and Supplement 2). Two upper bands (*ca*. 40 and 55 kDa) were found to contain only one ribosomal protein each, namely, uL4 and eL6, whereas the lower band (*ca*. 20 kDa) included two proteins, uL13 and eL15. Thus, proteins uL4 (RPL4), eL6 (RPL6), and uL13/eL15 (RPL13A/RPL15) are the main targets of modification by PARP1 among 60S RPs. Since the positions of the bands of 60S RPs PARylated by PARP2 are similar to those with PARP1 (Figure 3a, b, lanes 12), one can assume that both PARPs modify the same 60S proteins. Considering the lower yield of the HPF1-dependent PARylation of 40S RPs as compared to that of 60S ones (see above) and a difference between PARylation patterns of 40S RPs with PARP1 and PARP2, we decided not to perform here the identification of the labeled 40S RPs.

### Interplay between PARylation of RPs and histones

Finally, we examined what happens when both RPs and histones are present together in the mixtures under conditions of ADP-ribosylation with PARP1 or PARP2 in the presence of HPF1, which indeed acts as a co-factor during histones PARylation in the nucleus. For this purpose, we used either a linker histone H1 alone or core histones. We did not use in these experiments the nucleosomal particles due to possible interference of nucleosmal DNA as an activator in the process of PARylation of RP with free DNA applied in our experiments as well as regarding other difficulties in data interpretation. We used total RP from 40S or 60S subunits at 0.45 µM concentration and histones at the same or 5-fold higher concentrations. The results in Figure 4 show that histone H1 taken at the same concentration as total RP is PARylated much less effectively than RPs. At that, the histone does not significantly affect the RPs PARylation in the mixture, where the RPs and H1 are present in the equimolar concentration (Figure 4a and 4c). Somewhat inhibitory effect of H1 on the PARylation of RPs becomes detectable only when it is taken in a 5-fold excess to RPs (Figure 4b and 4d); surprisingly, H1 in this case remains unmodified. All this implies that linker histone H1 is a poor inhibitor of PARylation of both 60S and 40S RPs with both PARPs. At that, RPs themselves very effectively prevent H1 PARylation.

**Figure 4.**
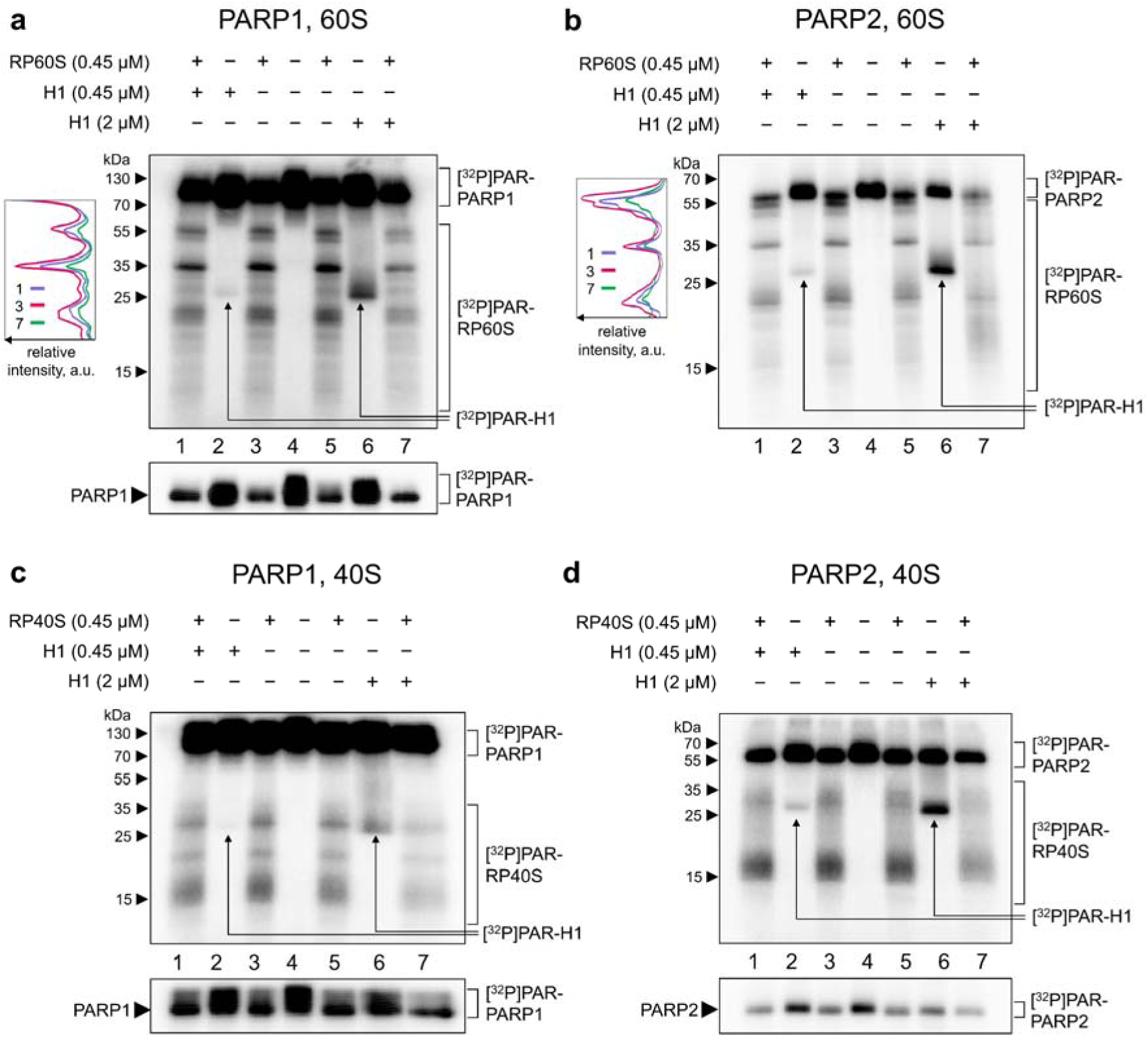
Effect of linker histone H1 on PARylation of ribosomal proteins with PARP1 and PARP2. Covalent binding of ^32^P-labelled ADP-ribose to 60S (**a, b**) or 40S (**c, d**) total ribosomal proteins in the presence of linker histone H1. Concentrations of the PARPs, dsDNA-activator, HPF1 and [^32^P]NAD^+^ were as in Figure 3; concentrations of ribosomal proteins and histone H1 are indicated above the panels where applied. Diagrams representing the distribution of the intensity among the bands are shown on the left of the autoradiograms (**a**) and (**b**). Autoradiograms of the gels after separation of the products by 15% SDS-PAGE. Parts of the autoradiographs with reduced exposure that contain only the PARP signals are given at the bottom.

Experiments with core histones similar to those described above with histone H1 lead to strikingly different results (Figure 5). These histones were PARylated by both PARPs more effectively, than RPs, and when taken in an equimolar 0.45 µM concentration with total RP, the core histones noticeably inhibited PARylation of the RPs and simultaneously underwent PARylation themselves. This was observed with both 40S and 60S RPs and with both PARPs. When taken in a 5-fold excess over RPs, the core histones completely inhibited their PARylation. Thus, these histones are strong competitors with both 40S and 60S RPs for PARP1/2.

**Figure 5.**
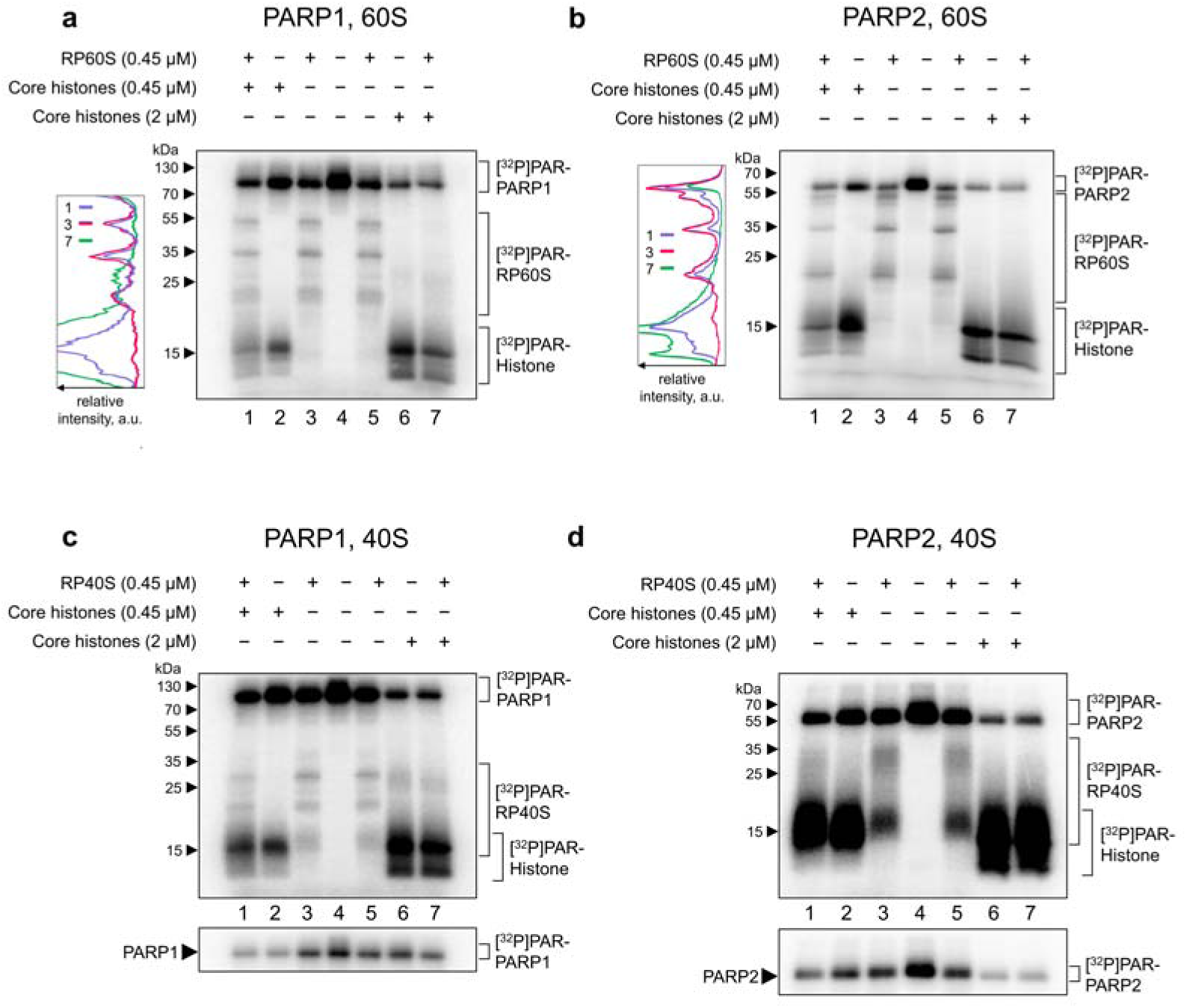
Interrelations between PARylation of ribosomal proteins and core histones with PARP1 and PAPR2. Covalent binding of ^32^P-labelled ADP-ribose to 60S (**a, b**) or 40S (**c, d**) total ribosomal proteins in the presence of core histones. Concentrations of the ribosomal proteins, PARPs, dsDNA-activator, HPF1 and [^32^P]NAD^+^ were as in Figure 4; concentrations of the core histones are indicated above the panels where applied. Diagrams representing the distribution of the intensity among the bands are shown on the left of the autoradiograms (**a**) and **(b**). Autoradiograms of the gels after separation of the products by 15% SDS-PAGE.

Our results are summarized in the scheme presented in Figure 6.

**Figure 6.**
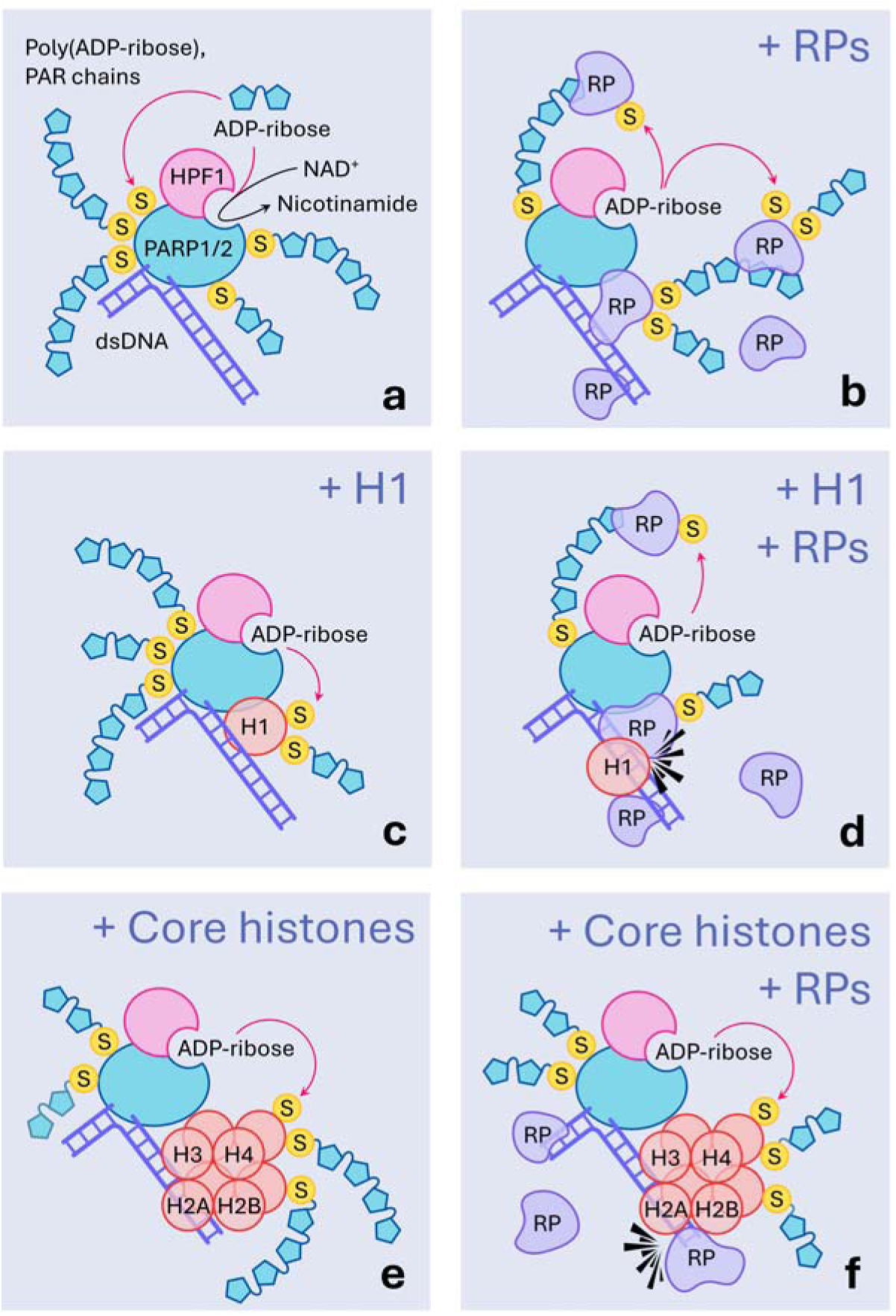
Scheme showing the putative involvement of RPs into the interactions of PARP1/2 with damaged DNA. (**a**) In the presence of HPF1, PARP1/2 modify themselves with short PAR chains at serine residues (S). (**b**) RPs, if present, bind to DNA and/or to the PAR chains (?) and undergo PARylation, which considerably reduces the PARP1/2 automodification. (**c**) Linker histone H1, when occurs near the complex of damaged DNA with PARP1/2 and HPF1, is also PARylated, although less effectively than RPs, and its PARylation has almost no effect the PARP1/2 automodification. (**d**) When RPs and histone H1 are present together, RPs prevent the binding of the latter to DNA or PAR chains near the PARPs and undergo PARylation. (**e**) Core histones are effectively PARylated in the vicinity of the complex of PARP1 (or PARP2) and HPF1 with damaged DNA, which reduces the level of the PARP automodification. (**f**) When RPs arrive to the above complex, they compete with core histones for PARP, apparently due to the competition for binding sites on the damaged DNA and/or PAR chains. If the core histones are in excess compared to RPs, they win the competition and the RPs remain unmodified.

## Discussion

Our results provide a response to the question arising from the studies of human PARylome [6-7, 20] concerning whether RPs are PARylated within assembled ribosomal subunits or outside them. We used here mature 40S and 60S subunits, and aware that their structures are not entirely the same as the pre-ribosomal particles present in nucleus, where they can encounter with PARP1 or PARP2. Nevertheless, the mature subunits, although being inexact models, seems applicable since it is known that almost all RPs are already assembled into the subunits in nucleus, and their final maturation in the cytosol does not change the set of RPs in the subunits (e.g., see [30-31]). We show that under near-physiological conditions (buffer, ionic strength and concentrations of RPs in the micromolar range (e.g., see Supplement 3 to the report [32]), RPs of both 60S and 40S ribosomal subunits can undergo PARylation with both PARP1 or PARP2 in ribosome-free state but not within the ribosomal subunits. This is consistent with the earlier suggestion that the sites of RPs PARylation are those that implicated in the ribosome biogenesis and inaccessible within the assembled ribosome because of intra-ribosomal protein-protein or protein-rRNA interactions [6].

PARylation of both 60S or 40S ribosome-free RPs with either PARP1 or PARP2 observed in this study completely depends on the presence of a DNA activator as expected for reaction of PAR synthesis catalysed with PARP1 and PARP2 [33] We obtained here the evidence for the implication of HPF1 in PARylation of RPs. The effect of this co-factor on the 60S RPs modification in general is as expected from the earlier studies with PARylation of histones and the PARP enzymes themselves [34]. In particular, the HPF1 switches the modification mainly to the serine residues, strongly increases the amount of ADP-ribose resides and shortens the length of the ADP-ribose chain attached to the target protein [18, 35]. Somewhat surprisingly, this was not all the same with the 40S RPs. In the latter case, the HPF1-induced switch to the Ser residues took place, but without a simultaneous increase of the number of ADP-ribose residues per a target protein. This may be due to fewer serine residues accessible for PARylation in 40S RPs than that in 60S ones. Analysis of the respective modification sites in RPs identified earlier in [20] (summarized as Supplement 3) shows that the total number of PARylated serines in 60S RPs (102) is indeed three-fold of that in 40S RPs (34). Thus, a lower output of 40S RPs PARylation as compared to that of 60S ones might account, at least in part, for the smaller number of PARylated residues in the 40S RPs. However, the other explanation of this difference can be also found.

Noteworthy, our results demonstrating that HPF1 does not increase the extent of PARylation of 40S RPs differ from the data on strong stimulating effect of the HPF1 on the PARylation of recombinant RPS3A (eS1) by PARP1 *in vitro* [7]. With 40S RPs, we indeed observed some radioactive bands in the region of 30-35 kDa, where RPS3A migrates (Figures 1 and 2), but the intensity of these bands does not increase in an HPF1-dependent manner. A possible explanation for this inconsistency could be different kinds of activated DNA used here and in the mentioned report as well as other conditions of the performed experiments [7].

PARylation of ribosome-free RPs simultaneously decreases the yield of auto-PARylation of the respective PARP (see e.g., Figure 2). This effect is observed with both kinds of RPs (40S and 60S) and both PARPs although the details vary in each case. The reason for the effect, at least in part, could be that RPs are competitive targets of PARylation with the enzymes and take away NAD^+^ available for PARP’s automodification.

Our results demonstrate that certain RPs are preferentially PARylated, which is especially clearly seen with 60S RPs in the presence of HPF1. In the latter case, the quantification of the results shown in Figures 1-3 enables an estimation that 70-90% of PAR residues attached to 60S RPs mainly concern three or four modified RPs, RPL4 (uL4), RPL6 (eL6) and uL13/eL15 (RPL13A/RPL15) of total 47 large subunit RPs. Currently, it is difficult to exactly explain why the PARPs are selectively attracted to these RPs as targets of trans-PARylation, and this effect might originate from several reasons. The first reason is a larger amount of Ser residues suitable for trans-PARylation by PARPs in the presence of HPF1. However, the former reason seems unlikely because some non PARylated 60S RPs have a number of the PARylatable Ser residues similar to those in RPL4 (uL4), RPL6 (eL6) and uL13/eL15 (RPL13A/RPL15) (see Supplement 3). Besides, the selective PARylation of definite 60S RPs might be provided (at least, in part) by their interaction with DNA and/or with PAR chains attached to PARPs as the result of their autoPARylation (Figure 6). This interaction is expectable because most RPs are positively charged while DNA and PAR are charged negatively, and differences in the interactions of specific RPs with DNA and/or PAR chain could, at least in part, result in preferential PARylation of the selected RPs. It is possible that RPs, when their level is high, interact with HPF1-PARP complex bound at the damaged DNA and under these conditions RPs become the main targets for trans-PARylation (such kind events have been described with YB1 protein as a partner of PARP1 [12, 36]). This could be a possible reason for the decrease in the PARylation level of PARPs and 60S RPs at high but not low concentrations of the 60S RPs (Figure 2a).

Remarkably, RPL6 (eL6), which is PARylated most preferentially among 60S RPs with both PARP1/2, has been reported earlier to play a role as a critical regulatory factor in DNA damage response [37]. This RP is recruited to DNA damage sites in a PARP-dependent manner and initiates a signaling cascade by facilitating the interaction of a mediator of DNA damage checkpoint 1 (MDC1) with phosphorylated H2AX histone (γH2AX), which eventually promotes the recruitment of a DNA repair protein 53BP1 and BRCA1 [37]. Although it remained unclear whether RPL6 undergoes PARylation itself in the course of the above events, our results support the latter possibility. In addition to RPL6, two more RPs, RPS19 (eS19) and RPL5 (uL18), are known to be recruited to DNA damage sites in a PARP-dependent manner; these RPs interact with Ku70 and histone H2A, and also with PAR chains [38]. We do not detect here a significant PARylation of RPL5, and therefore one can suggest that implication of this RP in DNA damage response might occur without covalent attachment of PAR chains or PARylation of RPL5 is not effective under our conditions. Currently, our results do not allow making any grounded suggestions concerning the implication of RPS19 since we did not set ourselves the goal to identify PARylated 40S RPs.

Examination of PARylation of RPs and various histones by PARP1 or PARP2 in the common mixture represent a simplified model of a situation that could be realized upon stress leading to DNA damage and arrest of ribosome biogenesis resulting in the accumulation of ribosome-free RPs in nucleus. Our results show that linker histone H1 is PARylated in the presence of HPF1 less effectively than RPs with both PARP1 and PARP2, and it is a poor competitor with them for the enzymes. Noteworthy, RPs PARylation is somewhat inhibited in the presence of 5-fold excess of H1, but at that, this histone did not undergo PARylation itself (Figure 4).

Considering all the above, we can suggest a following consequence of events with participation of RPs following genotoxic stress (Figure 7). DNA damage outside the nucleolus leads to the kinase ATM-dependent arrest of transcription by RNA polymerase I, which results in the stall of ribosome biogenesis [39]. In the damaged region, HPF1-dependent PARylation of linker histone H1 occurs [40]. Attachment of negatively charged PAR chains to H1 decreases its affinity to the DNA and thereby causes its dissociation, which follows by the activation of kinase ATM. The arrest of ribosome biogenesis results in an accumulation of ribosome-free RPs in the nucleus (for review, see e.g. [41]), and they can appear at the DNA damage site where they can interact with PAR synthesized with PARPs [42-43]. It has been shown that loading of RPL6 to DNA damage site is inhibited by olaparib. Therefore, this process is dependent on PAR synthesis, and it was suggested that RPL6 encounters histones [37]. Considering our results on the strong inhibitory action of RPs on histone H1 PARylation, one can assume that PARPs stop the H1 modification as soon as RPs arrive (Figure 6). Functional significance of the RPs’ action could be prevention an excessive PARylation of H1, which potentially might lead to the undesirable decompactization of chromatin far from the DNA damage site.

**Figure 7.**
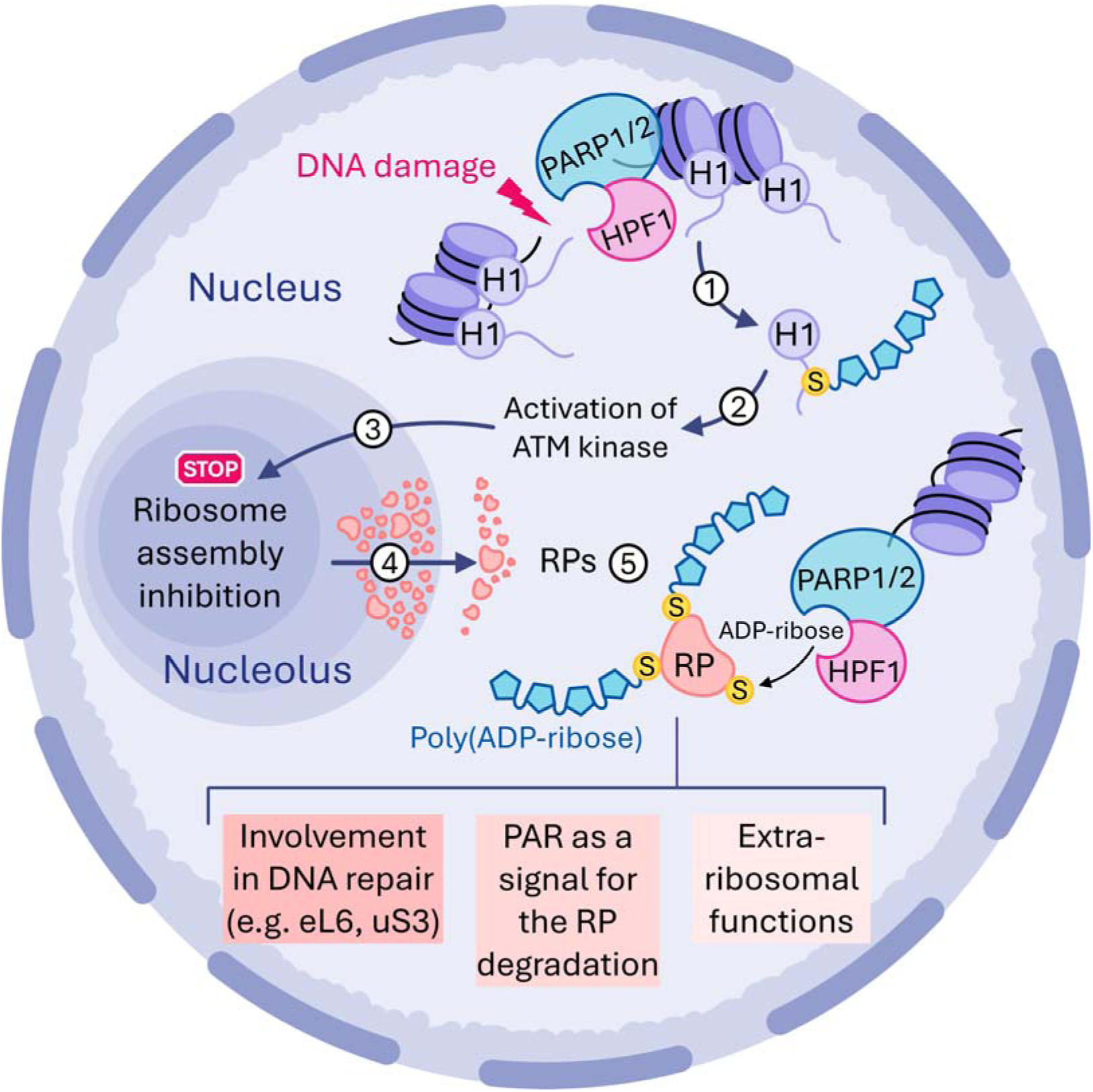
Hypothetic scheme of events leading to PARylation of RPs upon DNA damage in the nucleus and consequences of this post-translational modification. **(1)** DNA damage outside the nucleolus leads to the activation of PARP1/2 and HPF1-dependent PARylation of linker histone H1 in the damage region. Negatively charged PAR chains on the H1 decreases affinity of the latter to the DNA and results in the histone dissociation. (**2**) Dissociation of the linker histone results in the activation of ATM kinase. (**3**) The latter event leads to the arrest of transcription by RNA polymerase I, which results in the stall of ribosome biogenesis. **(4)** Arrest of the ribosome biogenesis leads to an accumulation of ribosome-free RPs in the nucleoplasm, where they can be PARylated. (**5**) PARylated RPs are involved in the DNA repair and/or perform their extra-ribosomal functions. Besides, PAR chains attached to the RPs can serve as a signal triggering degradation of the excessive RPs, which might be mediated, for instance, by the recently discovered ubiquitination of ADP-ribose [46] attached to the proteins.

What else could be functional assignments of PARylation of RPs? Considering that excessive free RPs accumulating in the nucleus as a consequence of the stall of ribosome biogenesis are toxic for the cell (e.g., form aggregates [44]), PARylation might be associated with pathways leading to degradation of RPs. In this respect, one can mention ubiquitin ligase HUWE1 that has been shown to ubiquitinate excessive ribosome-free RPs for their subsequent proteasomal degradation [44]. This enzyme contains a PAR binding WWE domain [45], which makes reasonable a suggestion on implication of RPs PARylation in their degradation. One can also assume that PARylation of RPs prevents their participation in the ribosomal subunits assembly, and their PARylation may be one of the responses to stress leading to the arrest of ribosome biogenesis (Figure 7).

Finally, PARylation of ribosome-free RPs might relate to a regulation of their extra-ribosomal functions (Figure 7). However, currently there is too scarce information for a reasonable discussion of such kind possibility, and further investigations might shed light on this issue.

## Supporting information

Scores and coverage rate of identified ribosomal proteins

Identification of the major modified 60S proteins by MALDI TOF mass spectrometry

Serine PARylation sites in RPs

## AUTHOR CONTRIBUTIONS STATEMENT

Conceptualization, O.I.L., D.M.G. and D.P.; methodology, A.S.K., K.N.N., M.M.K., M.O.P. and J.B.Z.; validation, A.S.K., K.N.N. and D.M.G.; formal analysis, A.S.K., K.N.N. and M.O.P.; investigation, A.S.K., K.N.N., M.M.K., M.O.P. and J.B.Z.; resources, O.I.L. and K.N.N.; writing—original draft preparation, A.S.K., K.N.N. and D.M.G.; writing—review and editing, O.I.L., A.A.M. and D.P.; visualization, A.S.K. and M.O.P.; supervision, O.I.L. and D.M.G.; project administration, O.I.L. and K.N.N.; funding acquisition, O.I.L. and K.N.N. All authors have read and agreed to the published version of the manuscript.

## References

1. Alemasova, E. E. & Lavrik, O. I. Poly(ADP-ribosyl)ation by PARP1: reaction mechanism and regulatory proteins. Nucleic Acids Res. 47, 3811–3827 (2019).

2. Challa, S., Khulpateea, B. R., Nandu, T., Camacho, C. V., Ryu, K. W., Chen, H., Peng, Y., Lea, J. S. & Kraus, W.L. Ribosome ADP-ribosylation inhibits translation and maintains proteostasis in cancers. Cell. 184, 4531–4546.e26 (2021).

3. Challa, S., Nandu, T., Kim, H. B., Gong, X., Renshaw, C. W., Li, W. C., Tan, X., Aljardali, M. W., Camacho, C. V., Chen, J. & Kraus, W. L. RACK1 MARylation regulates translation and stress granules in ovarian cancer cells. J Cell Biol. 224, e202401101 (2025).

4. Bai, P. Biology of Poly(ADP-Ribose) Polymerases: The Factotums of Cell Maintenance. Mol Cell. 58, 947–958 (2015).

5. Suskiewicz, M. J., Munnur, D., Strømland, Ø., Yang, J. C., Easton, L. E., Chatrin, C., Zhu, K., Baretić, D., Goffinont, S., Schuller, M., Wu, W. F., Elkins, J. M., Ahel, D., Sanyal, S., Neuhaus, D. & Ahel, I. Updated protein domain annotation of the PARP protein family sheds new light on biological function. Nucleic Acids Res. 51, 8217–8236 (2023).

6. Zhen, Y., Zhang, Y. & Yu, Y. A Cell-Line-Specific Atlas of PARP-Mediated Protein Asp/Glu-ADP-Ribosylation in Breast Cancer. Cell Rep. 21, 2326–2337 (2017).

7. Leslie Pedrioli, D. M., Leutert, M., Bilan, V., Nowak, K., Gunasekera, K., Ferrari, E., Imhof, R., Malmström, L. & Hottiger, M. O. Comprehensive ADP-ribosylome analysis identifies tyrosine as an ADP-ribose acceptor site. EMBO Rep. 19, e45310 (2018).

8. Schreiber, V., Amé, J. C., Dollé, P., Schultz, I., Rinaldi, B., Fraulob, V., Ménissier-de Murcia, J. & de Murcia, G. Poly(ADP-ribose) polymerase-2 (PARP-2) is required for efficient base excision DNA repair in association with PARP-1 and XRCC1. J Biol Chem. 277, 23028–36 (2002).

9. Kutuzov, M. M., Khodyreva, S. N., Amé, J. C., Ilina, E. S., Sukhanova, M. V., Schreiber, V. & Lavrik, O. I. Interaction of PARP-2 with DNA structures mimicking DNA repair intermediates and consequences on activity of base excision repair proteins. Biochimie. 95, 1208–15 (2013).

10. Szántó, M., Yélamos, J. & Bai, P. Specific and shared biological functions of PARP2 - is PARP2 really a lil’ brother of PARP1? Expert Rev Mol Med. 26, e13 (2024).

11. Fischbach, A., Krüger, A., Hampp, S., Assmann, G., Rank, L., Hufnagel, M., Stöckl, M. T., Fischer, J. M. F., Veith, S., Rossatti, P., Ganz, M., Ferrando-May, E., Hartwig, A., Hauser, K., Wiesmüller, L., Bürkle, A. & Mangerich, A. The C-terminal domain of p53 orchestrates the interplay between non-covalent and covalent poly(ADP-ribosyl)ation of p53 by PARP1. Nucleic Acids Res. 46, 804–822 (2018).

12. Naumenko, K. N., Sukhanova, M. V., Hamon, L., Kurgina, T. A., Alemasova, E. E., Kutuzov, M. M., Pastré, D. & Lavrik, O. I. Regulation of Poly(ADP-Ribose) Polymerase 1 Activity by Y-Box-Binding Protein 1. Biomolecules. 10, 1325 (2020).

13. Bonfiglio, J. J., Fontana, P., Zhang, Q., Colby, T., Gibbs-Seymour, I., Atanassov, I., Bartlett, E., Zaja, R., Ahel, I. & Matic, I. Serine ADP-Ribosylation Depends on HPF1. Mol Cell. 65, 932–940.e6 (2017).

14. Palazzo, L., Leidecker, O., Prokhorova, E., Dauben, H., Matic, I. & Ahel, I. Serine is the major residue for ADP-ribosylation upon DNA damage. Elife. 7, e34334 (2018).

15. Suskiewicz, M. J., Zobel, F., Ogden, T. E. H., Fontana, P., Ariza, A., Yang, J. C., Zhu, K., Bracken, L., Hawthorne, W. J., Ahel, D., Neuhaus, D. & Ahel, I. HPF1 completes the PARP active site for DNA damage-induced ADP-ribosylation. Nature. 579, 598–602 (2020).

16. Suskiewicz, M. J., Palazzo, L., Hughes, R. & Ahel, I. Progress and outlook in studying the substrate specificities of PARPs and related enzymes. FEBS J. 288, 2131–2142 (2021).

17. Kurgina, T. A., Moor, N. A., Kutuzov, M. M., Naumenko, K. N., Ukraintsev, A. A. & Lavrik, O. I. Dual function of HPF1 in the modulation of PARP1 and PARP2 activities. Commun Biol. 4, 1259 (2021).

18. Rudolph, J., Roberts, G., Muthurajan, U. M. & Luger, K. HPF1 and nucleosomes mediate a dramatic switch in activity of PARP1 from polymerase to hydrolase. Elife. 10, e65773 (2021).

19. Vyas, S., Matic, I., Uchima, L., Rood, J., Zaja, R., Hay, R. T., Ahel, I. & Chang, P. Family-wide analysis of poly(ADP-ribose) polymerase activity. Nat Commun. 5, 4426 (2014).

20. Hendriks, I. A., Larsen, S. C. & Nielsen, M. L. An Advanced Strategy for Comprehensive Profiling of ADP-ribosylation Sites Using Mass Spectrometry-based Proteomics. Mol Cell Proteomics. 18, 1010–1026 (2019).

21. Boamah, E. K., Kotova, E., Garabedian, M., Jarnik, M. & Tulin, A.V. Poly(ADP-Ribose) polymerase 1 (PARP-1) regulates ribosomal biogenesis in Drosophila nucleoli. PLoS Genet. 8, e1002442 (2012).

22. Kutuzov, M. M., Kurgina, T. A., Belousova, E. A., Khodyreva, S. N. & Lavrik, O. I. Optimization of nucleosome assembly from histones and model DNAs and estimation of the reconstitution efficiency. Biopolym Cell. 35, 91–98 (2019).

23. Amé, J. C., Kalisch, T., Dantzer, F. & Schreiber, V. Purification of recombinant poly(ADP-ribose) polymerases. Methods Mol Biol. 780, 135–52 (2011).

24. Matasova, N. B., Myltseva, S. V., Zenkova, M. A., Graifer, D. M., Vladimirov, S. N. & Karpova, G. G. Isolation of ribosomal subunits containing intact rRNA from human placenta: estimation of functional activity of 80S ribosomes. Anal Biochem. 198, 219–23 (1991).

25. Hardy, S. J., Kurland, C. G., Voynow, P. & Mora, G. The ribosomal proteins of Escherichia coli. I. Purification of the 30S ribosomal proteins. Biochemistry. 8, 2897-905 (1969).

26. Sharifulin, D. E., Grosheva, A. S., Bartuli, Y. S., Malygin, A. A., Meschaninova, M. I., Ven’yaminova, A. G., Stahl, J., Graifer, D. M. & Karpova, G. G. Molecular contacts of ribose-phosphate backbone of mRNA with human ribosome. Biochim Biophys Acta. 1849, 930–9 (2015).

27. Daniels, C. M., Ong, S. E. & Leung, A. K. Phosphoproteomic approach to characterize protein mono- and poly(ADP-ribosyl)ation sites from cells. J Proteome Res. 13, 3510–22 (2014).

28. Zhang, Y., Wang, J., Ding, M. & Yu, Y. Site-specific characterization of the Asp- and Glu-ADP-ribosylated proteome. Nat Methods. 10, 981–4 (2013).

29. Cervantes-Laurean, D., Jacobson, E. L. & Jacobson, M. K. Preparation of low molecular weight model conjugates for ADP-ribose linkages to protein. Methods Enzymol. 280, 275–87 (1997).

30. Sanghai, Z. A., Miller, L., Molloy, K. R., Barandun, J., Hunziker, M., Chaker-Margot, M., Wang, J., Chait, B. T. & Klinge, S. Modular assembly of the nucleolar pre-60S ribosomal subunit. Nature. 556, 126–129 (2018).

31. Hartwick, E. W. & Wimberly, B. T. Resolving Late-Stage Intermediates of Eukaryotic Ribosome Assembly. Structure. 25, 216–218 (2017).

32. Hein, M. Y., Hubner, N. C., Poser, I., Cox, J., Nagaraj, N., Toyoda, Y., Gak, I. A., Weisswange, I., Mansfeld, J., Buchholz, F., Hyman, A. A. & Mann, M. A human interactome in three quantitative dimensions organized by stoichiometries and abundances. Cell. 163, 712–23 (2015).

33. Langelier, M. F., Riccio, A. A., Pascal, J. M. PARP-2 and PARP-3 are selectively activated by 5’ phosphorylated DNA breaks through an allosteric regulatory mechanism shared with PARP-1. Nucleic Acids Res. 42, 7762–75 (2014).

34. Suskiewicz, M. J., Prokhorova, E., Rack, J. G. M. & Ahel, I. ADP-ribosylation from molecular mechanisms to therapeutic implications. Cell. 186, 4475–4495 (2023).

35. Kurgina, T. A., Moor, N. A., Kutuzov, M. M. & Lavrik, O. I. The HPF1-dependent histone PARylation catalyzed by PARP2 is specifically stimulated by an incised AP site-containing BER DNA intermediate. DNA Repair (Amst). 120, 103423 (2022).

36. Naumenko, K. N., Sukhanova, M. V., Hamon, L., Kurgina, T. A., Anarbaev, R. O., Mangerich, A., Pastré, D. & Lavrik, O. I. The C-Terminal Domain of Y-Box Binding Protein 1 Exhibits Structure-Specific Binding to Poly(ADP-Ribose), Which Regulates PARP1 Activity. Front Cell Dev Biol. 10, 831741 (2022).

37. Yang, C., Zang, W., Ji, Y., Li, T., Yang, Y. & Zheng, X. Ribosomal protein L6 (RPL6) is recruited to DNA damage sites in a poly(ADP-ribose) polymerase-dependent manner and regulates the DNA damage response. J Biol Chem. 294, 2827–2838 (2019).

38. DeCleene, N. F., Asik, E., Sanchez, A., Williams, C. L., Kabotyanski, E. B., Zhao, N., Chatterjee, N., Miller, K. M., Wang, Y. H. & Bertuch, A. A. RPS19 and RPL5, the most commonly mutated genes in Diamond Blackfan anemia, impact DNA double-strand break repair. bioRxiv [Preprint]. 2024.10.10.617668 (2024).

39. Lindström, M. S., Jurada, D., Bursac, S., Orsolic, I., Bartek, J. & Volarevic, S. Nucleolus as an emerging hub in maintenance of genome stability and cancer pathogenesis. Oncogene. 37, 2351–2366 (2018).

40. Li, Z., Li, Y., Tang, M., Peng, B., Lu, X., Yang, Q., Zhu, Q., Hou, T., Li, M., Liu, C., Wang, L., Xu, X., Zhao, Y., Wang, H., Yang, Y. & Zhu, W. G. Destabilization of linker histone H1.2 is essential for ATM activation and DNA damage repair. Cell Res. 28, 756–770 (2018).

41. Boulon, S., Westman, B. J., Hutten, S., Boisvert, F. M. & Lamond, A. I. The nucleolus under stress. Mol Cell. 40, 216–27 (2010).

42. Gagné, J. P., Isabelle, M., Lo, K. S., Bourassa, S., Hendzel, M. J., Dawson, V. L., Dawson, T. M., & Poirier, G. G. Proteome-wide identification of poly(ADP-ribose) binding proteins and poly(ADP-ribose)-associated protein complexes. Nucleic acids res. 36, 6959–6976 (2008).

43. Gagné, J. P., Pic, E., Isabelle, M., Krietsch, J., Ethier, C., Paquet, E., Kelly, I., Boutin, M., Moon, K. M., Foster, L. J., & Poirier, G. G. Quantitative proteomics profiling of the poly(ADP-ribose)-related response to genotoxic stress. Nucleic acids res. 40, 7788–7805 (2012).

44. Sung, M. K., Porras-Yakushi, T. R., Reitsma, J. M., Huber, F. M., Sweredoski, M. J., Hoelz, A., Hess, S. & Deshaies, R. J. A conserved quality-control pathway that mediates degradation of unassembled ribosomal proteins. Elife. 5, e19105 (2016).

45. Krietsch, J., Rouleau, M., Pic, É., Ethier, C., Dawson, T. M., Dawson, V. L., Masson, J. Y., Poirier, G. G. & Gagné, J. P. Reprogramming cellular events by poly(ADP-ribose)-binding proteins. Mol Aspects Med. 34, 1066–87 (2013).

46. Kloet, M. S., Chatrin, C., Mukhopadhyay, R., van Tol, B. D. M., Smith, R., Rotman, S. A., Tjokrodirijo, R. T. N., Zhu, K., Gorelik, A., Maginn, L., Elliott, P. R., van Veelen, P. A., Ahel, D., Ahel, I. & van der Heden van Noort, G. J. Identification of RNF114 as ADPr-Ub reader through non-hydrolysable ubiquitinated ADP-ribose. Nat Commun. 16, 6319 (2025).

