## Supplementary material for "Interplay between human ribosomal proteins, PARP1, PARP2, HPF1 and histones": Identification of the major modified 60S proteins by MALDI TOF mass spectrometry

**Change in the intensity of radioactive signals of major [^32^P]ADP-ribosylated 60S RPs after the PARG treatment.**

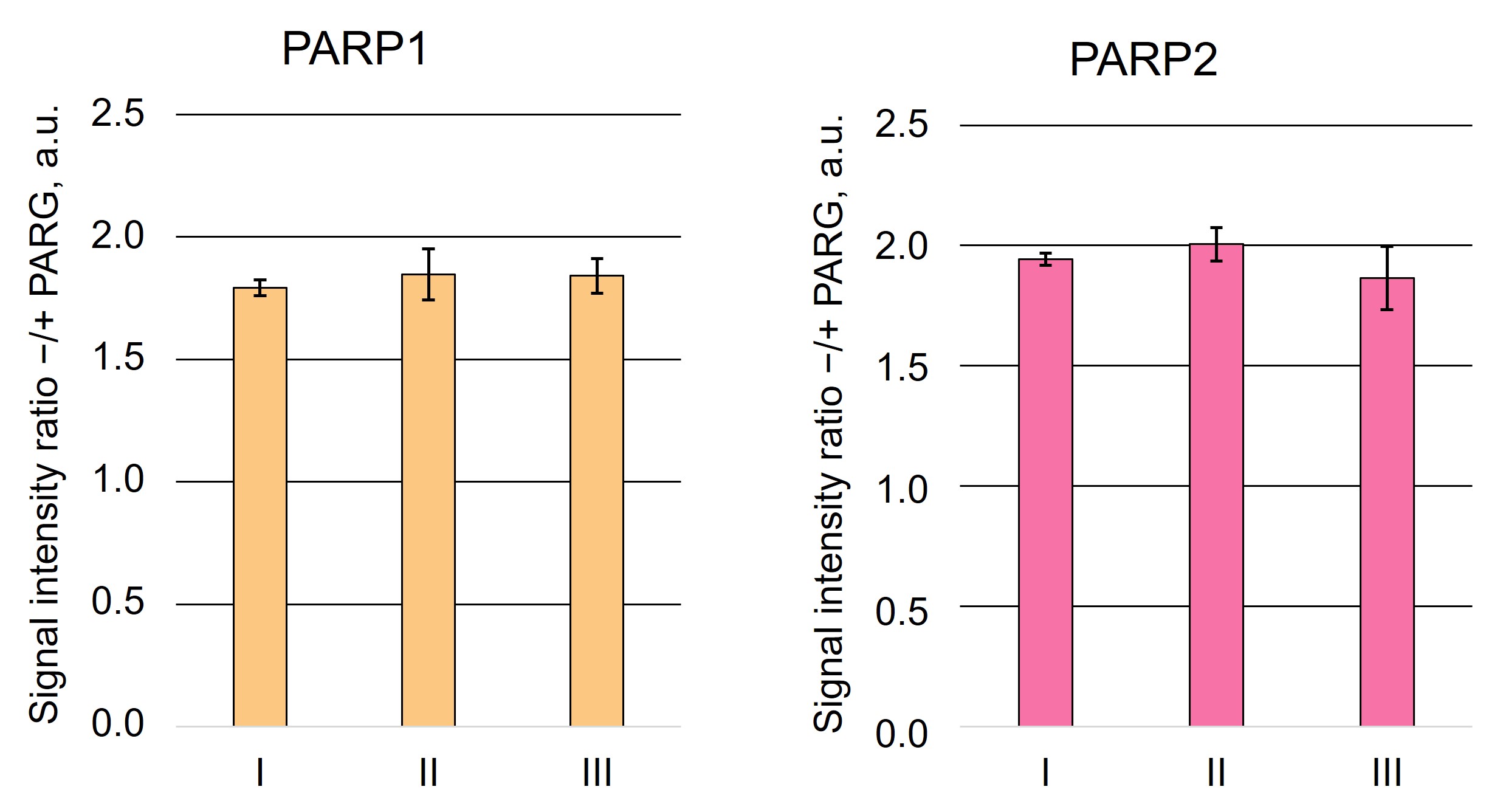

**Figure S1**. Histograms reflecting the ratio of the intensities of radioactive signals of the PARylated 60S RPs (major bands I, II, and III) before incubation with PARG to those after the incubation with PARG.The results are presented as the mean of arbitrary units (a.u.) from three independent repeats of the experiment ± SD.

**Preparation of samples of ADP-ribosylated 60S ribosomal proteins for identification of the major modified proteins by MALDI TOF mass spectrometry**

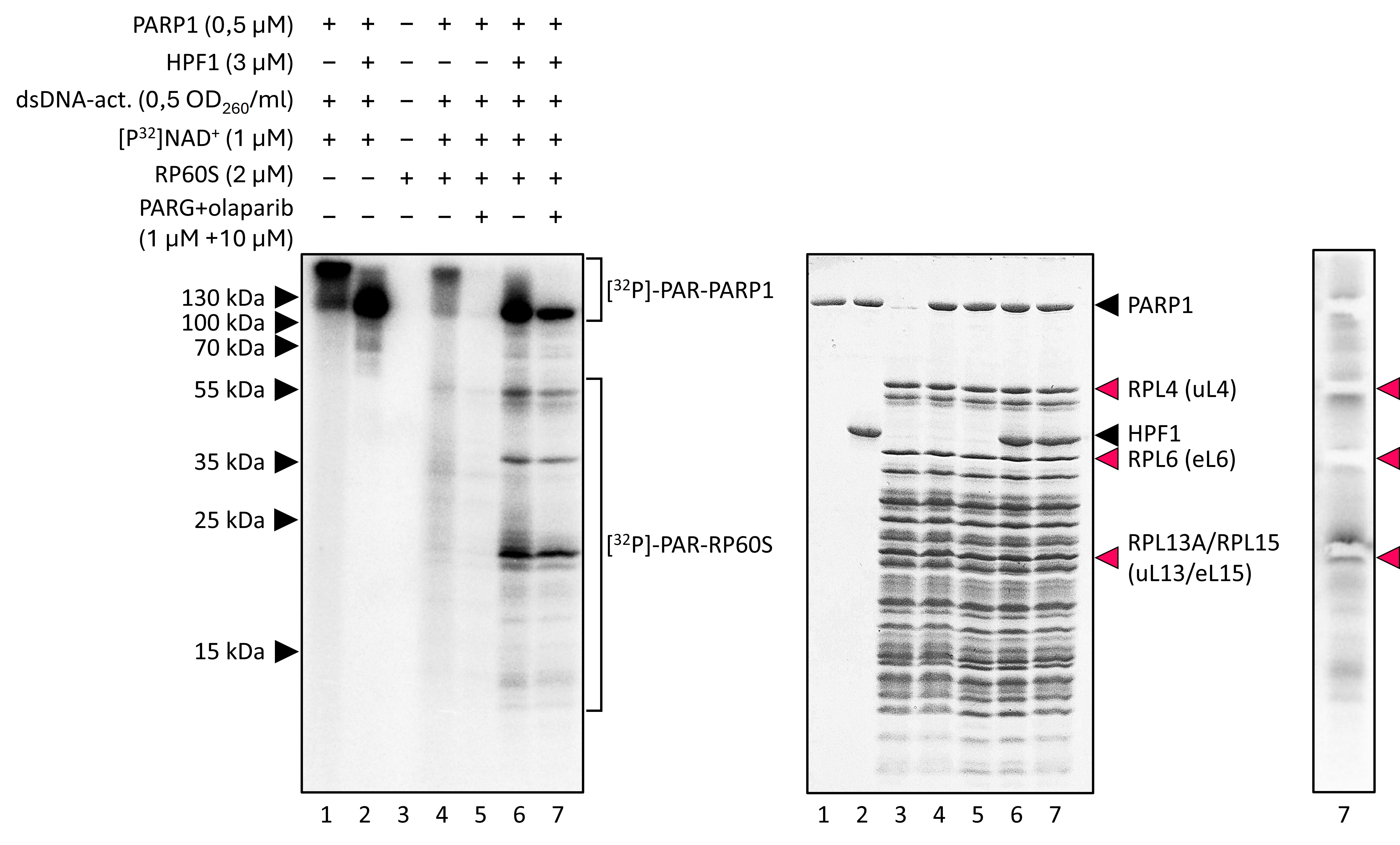
A	B	C

**Figure S2.** SDS-PAGE separation of RPs after their PARylation with PARP1, [^32^P]NAD^+^ for their subsequent identification by MALDI TOF mass spectrometry. **A**, autoradiogram; **B**, stained gel that has been subjected to autoradiography. Lane 7 was used for cutting pieces of the gel for the subsequent treatments to prepare samples for identification of RPs by MALDI TOF mass spectrometry. Bands to be cut off are shown at the stained gel (B). Panel **C** shows autoradiogram of the lane 7 after cutting off the main radioactive bands to ensure that cut off were indeed the desired bands. Lanes 1-6 are reference ones to control regularities of the PARylation of 60S RPs and autoPARylation of PARP1 observed in this study.

**Sample preparation for MALDI TOF**

Radioactive bands of interest were excised from the gels, cut into pieces of approximately 1 mm width, destained by washing with 50% CH_3_CN containing 25 mM NH_4_HCO_3_ and dried in vacuum. In-gel digestion with trypsin (sequencing grade modified, Promega) of proteins was carried out in the presence of ProteaseMAX Surfactant (Promega) that improves the protein digestion and peptide recovery from a gel. The digestion was performed by incubation of dried gel pieces with 20 μl of a solution containing 12 ng/μl of trypsin, 0.01% ProteaseMAX Surfactant and 50 mM NH_4_HCO_3_ at 0 °C for 30 min. Then 30 μl of 0.01% ProteaseMAX Surfactant in 50 mM NH_4_HCO_3_ was added, the mixture was incubated without agitation at 37°С (for the 1st replicate) or at 25 °С (for the 2nd replicate) overnight and then incubated at RT with agitation for 30 min. Gel pieces were separated from the solution of peptides by centrifugation for 2 min at 5,000 g. The experiment was performed in two replicates. In parallel, total 60S was resolved and coomassie-stained stained bands, whose positions corresponded to those of radioactive bands of interest, were excised from the gel and treated as described above (these samples were referred as “unmodified sample”, while the samples obtained with PARylated RPs are referred as “modified”).

MALDI TOF data obtained as the result of analysis of the peptides obtained and purified as described, are summarized in Supplementary Tables 1-16.

Below, the results of MALDI TOF analysis are presented as lists of coverage for each RP according to the data of Supplementary Tables 1-16.

**List of protein coverages.**

**uL4**

1^st^ replicate, unmodified sample

| **1** | MACARPLISV | YSEKGESSGK | **NVTLPAVFKA** | **PIRPDIVNFV** | **HTNLR**KNNR**Q** |
| --- | --- | --- | --- | --- | --- |
| **51** | **PYAVSELAGH** | **QTSAESWGTG** | **R**AVARIPRVR | GGGTHR**SGQG** | **AFGNMCR**GGR |
| **101** | MFAPTKTWRR | **WHRR**VNTTQK | RYAICSALAA | SALPALVMSK | GHR**IEEVPEL** |
| **151** | **PLVVEDK**VEG | YKKTKEAVLL | LKKLK**AWNDI** | **K**K**VYASQR**MR | AGKGKMRNRR |
| **201** | RIQR**RGPCII** | **YNEDNGIIKA** | **FRNIPGITLL** | **NVSK**LNILK**L** | **APGGHVGRFC** |
| **251** | **IWTESAFR**K**L** | **DELYGTWRK**A | ASLKSNYNLP | MHK**MINTDLS** | **RILKSPEIQR** |
| **301** | ALRAPRKKIH | RRVLKKNPLK | NLRIMLK**LNP** | **YAK**TMR**RNTI** | **LR**QARNHKLR |
| **351** | VDKAAAAAAA | LQAKSDEKAA | VAGKKPVVGK | KGKKAAVGVK | KQKKPLVGKK |
| **401** | AAATKKPAPE | KKPAEKKPTT | EEKKPAA |  |  |

1^st^ replicate, treated sample

| **1** | MACARPLISV | YSEKGESSGK | **NVTLPAVFKA** | **PIRPDIVNFV** | **HTNLR**KNNR**Q** |
| --- | --- | --- | --- | --- | --- |
| **51** | **PYAVSELAGH** | **QTSAESWGTG** | **R**AVARIPRVR | GGGTHR**SGQG** | **AFGNMCR**GGR |
| **101** | MFAPTKTWRR | **WHRR**VNTTQK | RYAICSALAA | SALPALVMSK | GHRIEEVPEL |
| **151** | PLVVEDK**VEG** | **YKK**TKEAVLL | LKKLK**AWNDI** | **K**KVYASQRMR | AGKGKMRNRR |
| **201** | RIQRRGPCII | YNEDNGIIKA | FRNIPGITLL | NVSKLNILK**L** | **APGGHVGRFC** |
| **251** | **IWTESAFR**K**L** | **DELYGTWRK**A | ASLK**SNYNLP** | **MHKMINTDLS** | **RILKSPEIQR** |
| **301** | ALRAPRKKIH | RRVLKKNPLK | NLRIMLKLNP | YAKTMR**RNTI** | **LR**QARNHKLR |
| **351** | VDK**AAAAAAA** | **LQAK**SDEKAA | VAGKKPVVGK | KGKKAAVGVK | KQKKPLVGKK |
| **401** | AAATKKPAPE | KKPAEKKPTT | EEKKPAA |  |  |

2^nd^ replicate, unmodified sample

| **1** | MACARPLISV | YSEKGESSGK | NVTLPAVFK**A** | **PIRPDIVNFV** | **HTNLR**KNNRQ |
| --- | --- | --- | --- | --- | --- |
| **51** | PYAVSELAGH | QTSAESWGTG | RAVARIPRVR | GGGTHR**SGQG** | **AFGNMCR**GGR |
| **101** | MFAPTKTWRR | WHRRVNTTQK | RYAICSALAA | SALPALVMSK | GHRIEEVPEL |
| **151** | PLVVEDKVEG | YKKTKEAVLL | LKKLK**AWNDI** | **K**KVYASQRMR | AGKGKMRNRR |
| **201** | RIQR**RGPCII** | **YNEDNGIIK**A | FRNIPGITLL | NVSKLNILK**L** | **APGGHVGRFC** |
| **251** | **IWTESAFR**K**L** | **DELYGTWRK**A | ASLKSNYNLP | MHKMINTDLS | R**ILKSPEIQR** |
| **301** | ALRAPRKKIH | RRVLKKNPLK | NLRIMLKLNP | YAKTMRR**NTI** | **LR**QARNHKLR |
| **351** | VDKAAAAAAA | LQAKSDEKAA | VAGKKPVVGK | KGKKAAVGVK | KQKKPLVGKK |
| **401** | AAATKKPAPE | KKPAEKKPTT | EEKKPAA |  |  |

2^nd^ replicate, treated sample

| **1** | MACARPLISV | YSEKGESSGK | NVTLPAVFK**A** | **PIRPDIVNFV** | **HTNLR**KNNR**Q** |
| --- | --- | --- | --- | --- | --- |
| **51** | **PYAVSELAGH** | **QTSAESWGTG** | **R**AVARIPRVR | GGGTHRSGQG | AFGNMCRGGR |
| **101** | MFAPTKTWRR | WHRRVNTTQK | RYAICSALAA | SALPALVMSK | GHRIEEVPEL |
| **151** | PLVVEDKVEG | YKKTKEAVLL | LKKLKAWNDI | KKVYASQRMR | AGKGKMRNRR |
| **201** | RIQRRGPCII | YNEDNGIIKA | FRNIPGITLL | NVSKLNILK**L** | **APGGHVGRFC** |
| **251** | **IWTESAFR**K**L** | **DELYGTWRK**A | ASLK**SNYNLP** | **MHK**MINTDLS | RILK**SPEIQR** |
| **301** | ALRAPRKKIH | RRVLKKNPLK | NLRIMLKLNP | YAKTMRRNTI | LRQARNHKLR |
| **351** | VDKAAAAAAA | LQAKSDEKAA | VAGKKPVVGK | KGKKAAVGVK | KQKKPLVGKK |
| **401** | AAATKKPAPE | KKPAEKKPTT | EEKKPAA |  |  |

**eL6**

1^st^ replicate, unmodified sample

| **1** | MAGEKVEKPD | TKEKKPEAKK | VDAGGKVKKG | NLKAKKPKKG | KPHCSR**NPVL** |
| --- | --- | --- | --- | --- | --- |
| **51** | **VR**GIGRYSR**S** | **AMYSRK**AMYK | RKYSAAKSKV | EKKKKEKVLA | TVTKPVGGDK |
| **101** | NGGTRVVKLR | KMPR**YYPTED** | **VPRK**LLSHGK | **KPFSQHVR**KL | R**ASITPGTIL** |
| **151** | **IILTGR**HRGK | RVVFLK**QLAS** | **GLLLVTGPLV** | **LNRVPLRR**TH | QKFVIATSTK |
| **201** | IDISNVKIPK | **HLTDAYFK**KK | KLRKPR**HQEG** | **EIFDTEKEKY** | **EITEQRK**IDQ |
| **251** | KAVDSQILPK | IK**AIPQLQGY** | **LR**SVFALTNG | IYPHKLVF |  |

1^st^ replicate, treated sample

| **1** | MAGEKVEKPD | TKEKKPEAKK | VDAGGKVKKG | NLKAKKPKKG | KPHCSR**NPVL** |
| --- | --- | --- | --- | --- | --- |
| **51** | **VR**GIGRYSR**S** | **AMYSRK**AMYK | RKYSAAKSKV | EKKKKEKVLA | TVTKPVGGDK |
| **101** | NGGTRVVKLR | KMPR**YYPTED** | **VPR**KLLSHGK | **KPFSQHVR**KL | R**ASITPGTIL** |
| **151** | **IILTGR**HRGK | RVVFLK**QLAS** | **GLLLVTGPLV** | **LNR**VPLRRTH | QKFVIATSTK |
| **201** | IDISNVKIPK | HLTDAYFKKK | KLRKPR**HQEG** | **EIFDTEKEKY** | **EITEQR**KIDQ |
| **251** | KAVDSQILPK | IK**AIPQLQGY** | **LR**SVFALTNG | IYPHKLVF |  |

2^nd^ replicate, unmodified sample

| **1** | MAGEKVEKPD | TKEKKPEAKK | VDAGGKVKKG | NLKAKKPKKG | KPHCSR**NPVL** |
| --- | --- | --- | --- | --- | --- |
| **51** | **VR**GIGRYSR**S** | **AMYSRKAMYK** | **RKYSAAK**SKV | EKKKKEKVLA | TVTKPVGGDK |
| **101** | NGGTRVVKLR | KMPR**YYPTED** | **VPR**KLLSHGK | **KPFSQHVR**KL | R**ASITPGTIL** |
| **151** | **IILTGR**HRGK | RVVFLK**QLAS** | **GLLLVTGPLV** | **LNR**VPLRRTH | QKFVIATSTK |
| **201** | IDISNVKIPK | HLTDAYFKKK | KLRKPR**HQEG** | **EIFDTEKEKY** | **EITEQR**KIDQ |
| **251** | KAVDSQILPK | IK**AIPQLQGY** | **LR**SVFALTNG | IYPHKLVF |  |

2^nd^ replicate, treated sample

| **1** | **MAGEKVEKPD** | **TK**EKKPEAKK | VDAGGKVKKG | NLKAKKPKKG | KPHCSR**NPVL** |
| --- | --- | --- | --- | --- | --- |
| **51** | **VR**GIGRYSR**S** | **AMYSRK**AMYK | RKYSAAKSKV | EKKKKEKVLA | TVTKPVGGDK |
| **101** | NGGTRVVKLR | KMPR**YYPTED** | **VPR**KLLSHGK | **KPFSQHVR**KL | R**ASITPGTIL** |
| **151** | **IILTGR**HRGK | RVVFLK**QLAS** | **GLLLVTGPLV** | **LNRVPLRR**TH | QK**FVIATSTK** |
| **201** | IDISNVKIPK | HLTDAYFKKK | KLRKPRHQEG | EIFDTEK**EKY** | **EITEQR**KIDQ |
| **251** | KAVDSQILPK | IK**AIPQLQGY** | **LR**SVFALTNG | IYPHKLVF |  |

**uL13**

1^st^ replicate, unmodified sample

| **1** | MAEVQVLVLD | GR**GHLLGR**LA | AIVAK**QVLLG** | **RKVVVVRCEG** | **INISGNFYR**N |
| --- | --- | --- | --- | --- | --- |
| **51** | KLK**YLAFLR**K | RMNTNPSR**GP** | **YHFR**APSR**IF** | **WR**TVRGMLPH | KTK**RGQAALD** |
| **101** | **R**LKVFDGIPP | PYDKKKRMVV | PAALKVVRLK | PTR**KFAYLGR** | **LAHEVGWKYQ** |
| **151** | **AVTATLEEK**R | KEKAKIHYRK | KKQLMRLRKQ | AEKNVEKKID | KYTEVLKTHG |
| **201** | LLV |  |  |  |  |

1^st^ replicate, treated sample

| **1** | MAEVQVLVLD | GR**GHLLGRLA** | **AIVAK**QVLLG | RKVVVVR**CEG** | **INISGNFYR**N |
| --- | --- | --- | --- | --- | --- |
| **51** | KLK**YLAFLR**K | RMNTNPSR**GP** | **YHFR**APSR**IF** | **WR**TVRGMLPH | KTKRGQAALD |
| **101** | RLKVFDGIPP | PYDKKKRMVV | PAALKVVRLK | PTR**KFAYLGR** | LAHEVGWK**YQ** |
| **151** | **AVTATLEEK**R | KEKAKIHYRK | KKQLMRLRKQ | AEKNVEKKID | KYTEVLKTHG |
| **201** | LLV |  |  |  |  |

2^nd^ replicate, unmodified sample

| **1** | MAEVQVLVLD | GR**GHLLGR**LA | AIVAK**QVLLG** | **R**KVVVVR**CEG** | **INISGNFYR**N |
| --- | --- | --- | --- | --- | --- |
| **51** | K**LKYLAFLR**K | RMNTNPSR**GP** | **YHFR**APSR**IF** | **WR**TVRGMLPH | KTKRGQAALD |
| **101** | RLK**VFDGIPP** | **PYDK**KKRMVV | PAALKVVRLK | PTR**KFAYLGR** | LAHEVGWK**YQ** |
| **151** | **AVTATLEEK**R | KEKAKIHYRK | KKQLMRLRKQ | AEKNVEKKID | KYTEVLKTHG |
| **201** | LLV |  |  |  |  |

2^nd^ replicate, treated sample

| **1** | MAEVQVLVLD | GR**GHLLGR**LA | AIVAK**QVLLG** | **R**KVVVVR**CEG** | **INISGNFYR**N |
| --- | --- | --- | --- | --- | --- |
| **51** | K**LKYLAFLR**K | RMNTNPSR**GP** | **YHFR**APSR**IF** | **WR**TVRGMLPH | KTK**RGQAALD** |
| **101** | **R**LK**VFDGIPP** | **PYDK**KKRMVV | PAALKVVRLK | PTR**KFAYLGR** | LAHEVGWK**YQ** |
| **151** | **AVTATLEEK**R | KEKAKIHYRK | KKQLMRLRKQ | AEKNVEKKID | KYTEVLKTHG |
| **201** | LLV |  |  |  |  |

**eL15**

1^st^ replicate, unmodified sample

| **1** | MGAYK**YIQEL** | **WR**KK**QSDVMR** | FLLRVR**CWQY** | **RQLSALHR**AP | RPTRPDKAR**R** |
| --- | --- | --- | --- | --- | --- |
| **51** | **LGYK**AK**QGYV** | **IYR**IRVRRGG | R**KRPVPKGAT** | **YGKPVHHGVN** | **QLK**FAR**SLQS** |
| **101** | **VAEER**AGRHC | GALRVLNSYW | VGEDSTYK**FF** | **EVILIDPFHK** | AIR**RNPDTQW** |
| **151** | **ITKPVHK**HRE | MR**GLTSAGR**K | SRGLGKGHK**F** | **HHTIGGSR**RA | AWRRR**NTLQL** |
| **201** | **HR**YR |  |  |  |  |

1^st^ replicate, treated sample

| **1** | MGAYK**YIQEL** | **WR**KK**QSDVMR** | FLLRVR**CWQY** | **RQLSALHR**AP | RPTRPDKARR |
| --- | --- | --- | --- | --- | --- |
| **51** | LGYKAK**QGYV** | **IYR**IRVRRGG | RKRPVPKGAT | YGKPVHHGVN | QLKFAR**SLQS** |
| **101** | **VAEER**AGRHC | GALRVLNSYW | VGEDSTYKFF | EVILIDPFHK | AIR**RNPDTQW** |
| **151** | **ITKPVHK**HRE | MRGLTSAGRK | SRGLGKGHK**F** | **HHTIGGSR**RA | AWRRR**NTLQL** |
| **201** | **HR**YR |  |  |  |  |

2^nd^ replicate, unmodified sample

| **1** | MGAYK**YIQEL** | **WR**KK**QSDVMR** | FLLRVR**CWQY** | **RQLSALHR**AP | RPTRPDKAR**R** |
| --- | --- | --- | --- | --- | --- |
| **51** | **LGYKAKQGYV** | **IYR**IRVRRGG | RKRPVPK**GAT** | **YGKPVHHGVN** | **QLK**FAR**SLQS** |
| **101** | **VAEER**AGRHC | GALR**VLNSYW** | **VGEDSTYKFF** | **EVILIDPFHK** | AIR**RNPDTQW** |
| **151** | **ITKPVHK**HRE | MR**GLTSAGR**K | SRGLGKGHK**F** | **HHTIGGSR**RA | AWRRR**NTLQL** |
| **201** | **HR**YR |  |  |  |  |

2^nd^ replicate, treated sample

| **1** | MGAYK**YIQEL** | **WR**KK**QSDVMR** | FLLRVR**CWQY** | **RQLSALHR**AP | RPTRPDKAR**R** |
| --- | --- | --- | --- | --- | --- |
| **51** | **LGYK**AK**QGYV** | **IYR**IRVRRGG | RKRPVPK**GAT** | **YGKPVHHGVN** | **QLK**FAR**SLQS** |
| **101** | **VAEER**AGRHC | GALR**VLNSYW** | **VGEDSTYKFF** | **EVILIDPFHK** | AIR**RNPDTQW** |
| **151** | **ITKPVHK**HRE | MR**GLTSAGR**K | SRGLGKGHK**F** | **HHTIGGSR**RA | AWRRR**NTLQL** |
| **201** | **HR**YR |  |  |  |  |
